# Metabotropic Glutamatergic Signaling Adaptively Controls Trans-thalamic Communication

**DOI:** 10.64898/2026.07.25.740691

**Authors:** Alan Lai, Xiao-Jing Wang

## Abstract

Trans-thalamic circuits link cortical regions through higher-order sensory (HOs) thalamic nuclei to support cognition. These circuits are rich with metabotropic glutamate receptors (mGluRs), whose seconds-long dynamics conflict with the thalamus’s traditional relay role, leaving their function unknown. Here, we present a model of group I mGluR-mediated suppression of K^+^ leak currents in single neurons. Including mGluRs in a trans-thalamic network of spiking neurons revealed distinct modulatory roles for thalamocortical and corticothalamic mGluRs. In pyramidal neurons, thalamocortical mGluRs strengthen somato-dendritic coupling and dendritic plateau potentials, boosting feedforward propagation of sensory information to enhance decision-making in a detection task. In HOs thalamic neurons, corticothalamic mGluRs temporally integrate inputs to shift thalamic firing from burst to tonic, a prediction supported by pulvinar recordings in behaving animals. This effect increases detection of a novel stimulus (“wake-up call”) while attenuating detection of repeated stimuli. Therefore, mGluRs enable the HOs thalamus to dynamically modulate cortical communication.

## 1 Introduction

Conventional models of information processing in the brain are typically organized along a cortical hierarchy. However, it has become increasingly clear that higher-order sensory (HOs) thalamic nuclei also play a crucial role in cognitive function through trans-thalamic (cortico-thalamic-cortical) circuits that indirectly connect two cortical regions along a cortical hierarchy [1–5]. This trans-thalamic circuitry is organized into two distinct classes of glutamatergic connections, often referred to as ‘drivers’ and ‘modulators’ [6, 7]. Driver connections have long been theorized to rapidly relay information with high fidelity, while modulator connections adjust how target neurons respond to that information. This proposed distinction is supported by their robust differences in synaptic properties: drivers show paired-pulse depression and rely on fast ionotropic neurotransmission mediated by the *α*-amino-3-hydroxy-5-methyl-4-isoxazolepropionic acid (AMPA) and N-methyl-D-aspartate (NMDA) receptors. On the other hand, modulators show paired-pulse facilitation and can uniquely activate metabotropic glutamate receptors (mGluRs), a family of G-protein-coupled receptors that alter neuronal excitability for several seconds through intracellular second messenger signaling pathways [2, 8]. Although the physiology and anatomical properties of driver and modulatory connections have been extensively documented [9–13], it is not clear how their distinct features affect network dynamics and cognitive function. Particularly mysterious is the function of modulatory pathways in thalamic circuitry, whose engagement of mGluRs mediates slow synaptic signaling at timescales that present a direct challenge to the textbook view of the thalamus as a passive relay of information. Motivated by this, we seek a principled understanding of how mGluR signaling in trans-thalamic circuitry supports the HOs thalamus’s role in cognition.

Despite the ubiquitous expression of mGluRs throughout the central nervous system [14], their computational function in the brain remains underexplored. Previous computational work has focused mainly on fast ionotropic neurotransmission mediated by AMPA and NMDA receptors [15–17]. This is because mGluR-mediated currents are substantially smaller in peak amplitude than AMPAR-mediated currents, yet linger thousands of times longer [18]. Such slow, small-amplitude currents are typically ignored in single-neuron and network models concerned with millisecond-scale fast dynamics and information processing, with a few exceptions [19–21].

The present work focuses on postsynaptic group I mGluRs, which stand out due to their ability to substantially suppress 2-Pore K^+^ Leak Channels (K2P) [8, 22–24]. By modulating these key voltage-independent ion channels, group I mGluRs are well-positioned to shape the excitability and firing properties of single neurons and, in turn, the broader trans-thalamic network. We investigated how the molecular machinery underlying mGluR-mediated slow modulation in single neurons is linked to network-level dynamics. We begin by developing a biophysically constrained phenomenological model of group I mGluR-mediated EPSCs based on their ability to suppress K2P channels and incorporating this synaptic mechanism into models of increasing complexity: from a single-neuron to the entirety of the trans-thalamic circuit (see Fig. 1a for a complete circuit diagram). This multi-scale framework allows us to dissect how mGluR-mediated modulation shapes dynamics across levels of biological abstraction within the trans-thalamic circuit.

**Fig. 1.**
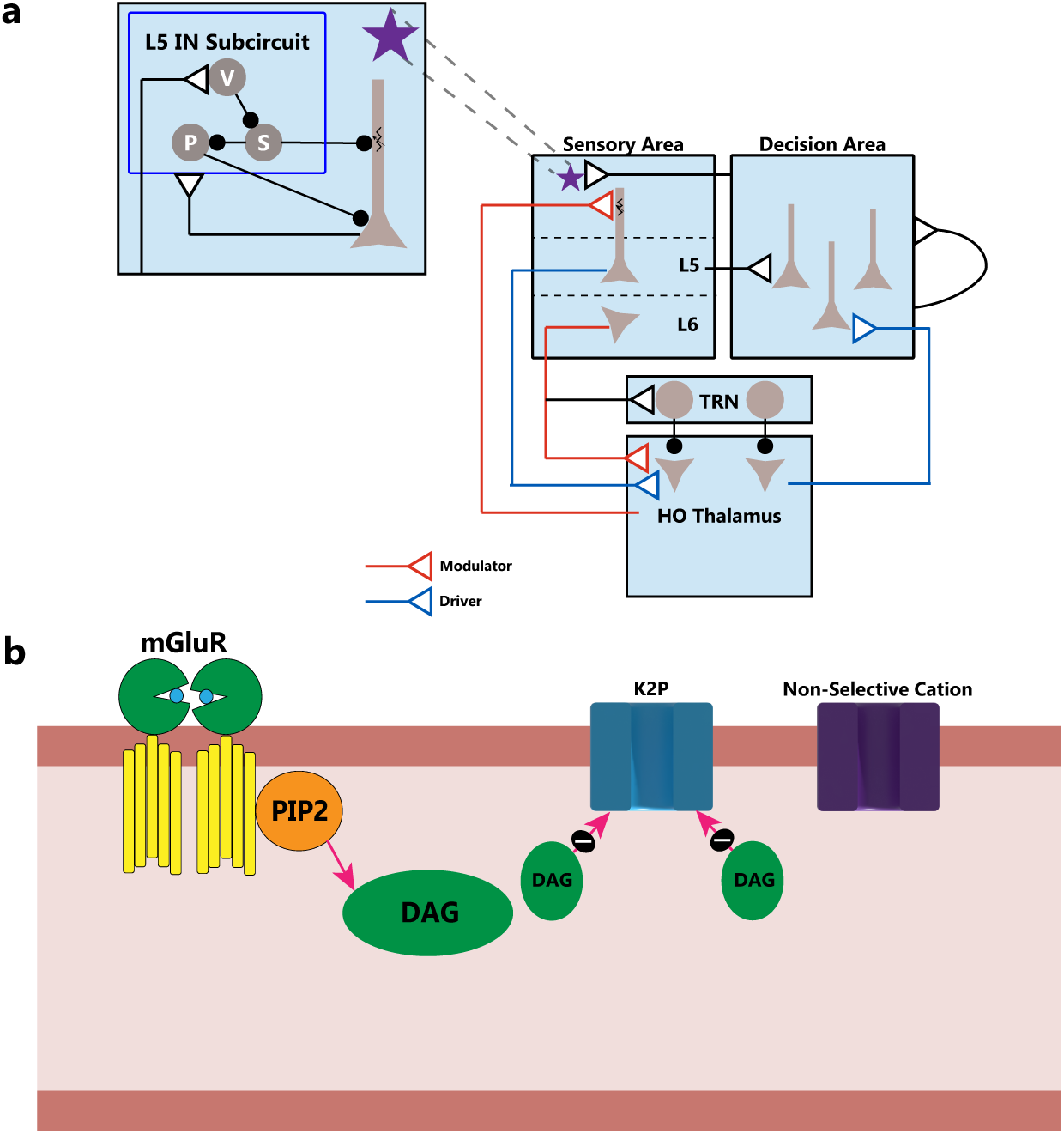
Model Schematics at different levels of biological abstraction. **(a.)** Trans-thalamic connections indirectly route information between cortical areas through the higher-order sensory (HOs) thalamus in parallel to direct cortical-cortical connections. Two main classes of excitatory glutamatergic connections (triangles) make up trans-thalamic circuitry: ‘Driver’ inputs (blue) are depressing synapses that do not engage mGluRs, while ‘modulator’ connections (red) are facilitating synapses that engage mGluRs. For simplicity, a third excitatory class (black) is modeled without short-term synaptic plasticity and engaging only ionotropic receptors. ‘Driver’ inputs carry feedforward information from sensory areas to the decision areas indirectly through HOs thalamus. Two ‘modulator’ pathways regulate the trans-thalamic circuit: Layer 6 corticothalamic (CT) projections modulate HOs thalamus, while thalamocortical (TC) feedback modulates the apical dendrites of Layer 5 pyramidal neurons (PNs). The thalamic reticular nucleus (TRN) forms a disynaptic inhibitory motif that inhibits (black circles) HOs thalamus. *Inset* : Layer 5 inhibitory neuron (IN) subcircuit. HOs thalamic neurons preferentially target VIP INs in sensory areas to disinhibit cortical dendrites, enabling dendritic plateau potentials. **(b.)** Phenomenological model of the mGluR EPSC. mGluR activation engages Gq-protein signaling to activate PLC*β* (not shown) to hydrolyze PIP2 and produce DAG, which inhibits K^+^ leak channels (K2P), resulting in a long-lasting depolarization.

Our analysis of thalamocortical (TC) modulation revealed that group I mGluR signaling can dynamically control the gain of cortical neurons in the sensory cortex. In a fully reconstructed multicompartmental model of a layer 5 (L5) pyramidal neuron (PN) in V1 [25, 26], TC feedback from the HOs thalamus onto the distal apical dendrites produced a prolonged mGluR-mediated depolarization that recruited voltage-dependent NMDA and high-threshold (L-type) Ca^2+^ channels essential for dendritic plateau potentials. In addition, this suppression of voltage-independent K2P channels enhanced somato-dendritic coupling by increasing dendritic membrane resistance and improving the electrotonic compactness of the apical dendrites. Together, we found that these effects introduced a dendritic nonlinearity in the neuronal input-output function that substantially increased the gain of L5 PNs.

We next asked how these modulatory effects on pyramidal neurons shape behavior. Previous theoretical work suggests that perceptual detection relies on a dynamical bifurcation in which a sufficiently strong feedforward drive triggers an all-or-none “ignition”, a transition from transient to sustained high-firing activity in decision-related cortical areas [27–29]. We reasoned that if TC activation of mGluRs regulates the gain of single neurons in the sensory cortex, it can also regulate the threshold at which this ignition occurs. To test this, we developed a trans-thalamic network of spiking neurons comprised of two cortical regions and one HOs thalamic nucleus capable of performing a detection task. In this network, we added our phenomenological model of mGluR-mediated EPSCs to modulatory connections and found that increasing the strength of mGluR modulation in TC modulatory connections boosted feedforward sensory propagation by increasing the gain of sensory neurons, lowering the perceptual threshold for detection.

We then study the role of the corticothalamic (CT) modulatory pathway, where we find that mGluRs can affect the firing mode of HOs thalamic neurons. In model HOs thalamic neurons, we find that model CT modulatory synapses can integrate feedforward CT input over seconds-long timescales through mGluR’s slow kinetics. With sufficient depolarization, thalamic neurons shift from burst to tonic firing via inactivation of low-threshold (T-type) Ca^2+^ channels. Because this depolarization decays over seconds, incoming excitatory input does not need to be confined to a single trial within an experimental session: CT mGluR-mediated depolarization can accumulate across successive behavioral trials, so that the thalamic firing mode reflects the recent trial history of CT input. This prediction is supported by the analysis of *in vivo* pulvinar recordings in awake-behaving mice, in which thalamic bursting significantly decreases across repeated trials of L6 CT stimulation.

Finally, we examine the behavioral role of this transition. Because TC modulation from the HOs thalamus is strongly facilitating, thalamic bursts evoke larger postsynaptic effects on cortical PNs than tonic spikes [30, 31]. Building on this, we found that in the trans-thalamic network of spiking neurons, the inclusion of mGluRs allowed CT modulator connections to accumulate CT depolarizing input in HOs thalamic neurons, causing thalamic neurons to fire tonically and reducing their modulatory effectiveness onto the sensory cortex. In doing so, CT modulation implements a “thalamic wake-up call” in which the network’s detection performance attenuates across repeated stimuli while novel input elicits a heightened response. Through this complementary organization, mGluR signaling acts distinctly along the two pathways: TC modulation amplifies the feedforward propagation of sensory input, while CT modulation regulates the strength of that amplification according to the recent history of CT input.

Altogether, our findings highlight that slow group I mGluR signaling transforms the thalamus from a passive relay into a dynamic, history-dependent regulator of cortical gain that governs how effectively sensory input is communicated across cortex.

## 2 Results

### 2.1 A Phenomenological Model for mGluR-mediated EPSCs

A key feature of mGluRs is their ability to modulate key ion channels over the course of several seconds through second-messenger signaling pathways. Given the complexity of the activated biochemical cascade and our goal of gaining a systems-level understanding of modulatory pathways in trans-thalamic circuitry, our first objective is to design a computationally tractable model of an essential function of group I mGluRs: suppression of K2P leak channels, which has been experimentally shown to be the underlying biophysical mechanism that depolarizes thalamic and cortical neurons following group I mGluR activation [22, 32].

Classically, group I mGluRs are coupled with G_q_ proteins which, upon their activation, stimulate phospholipase C*_β_* (PLC*β*) to hydrolyze phosphatidylinositol 4,5-bisphosphate (PIP2) into inositol 1,4,5-trisphosphate (IP3) and diacylglycerol (DAG) [14]. Although IP3 and DAG production mediate further downstream processes involving intracellular Ca^2+^ release and protein kinase C (PKC) activation, respectively, previous experimental work has identified DAG as a key molecule that can directly suppress K2P channels, independently of downstream pathways mediated by PKC [24, 33]. Consistent with this molecular characterization, a combination of transcriptomic and pharmacological analyses has suggested that heterogeneous expression of genes regulating DAG signaling across a neuronal population gives rise to a diversity of mGluR EPSC waveforms. Specifically, the expression of DAG kinases (DGKs), enzymes that terminate DAG signaling, is inversely correlated with the EPSC duration, while the expression of group I mGluRs and PLC*β* is positively correlated with the EPSC amplitude [34].

Based on these experimental results, we design a phenomenological model that captures both the long time-course of the mGluR EPSC as well as the EPSC’s dependence on DAG (see Fig. 1b for a schematic). In our model, we assume that the conductance of K2P leak channels can be multiplied by a factor *K*, which ranges over [*K_min_*, 1]; *K_min_ >* 0, and describes the proportion of K2P leak channels that are open. At rest, all K2P leak channels are active; however, after mGluR activation, DAG is produced and decreases *K* by a rate controlled by *β_K_*. When DAG disappears, the K2P channels will recover to their maximum conductance (*K* = 1) with a time constant of *τ_K_*. Mathematically, this is represented by the equation:

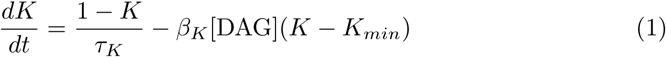

Then, to model intracellular DAG signaling, we use the following dynamical equation:

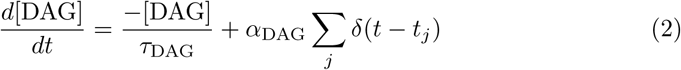

Here, a presynaptic spike at time *t_j_* produces a transient increase in postsynaptic intracellular DAG with an amplitude *α*_DAG_ that decays exponentially to zero with a time constant *τ*_DAG_. To generate a depolarization, we decompose the leak current into non-selective cation and K^+^ components and scale the K^+^ leak current by *K*.

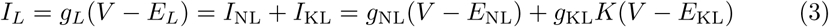

With

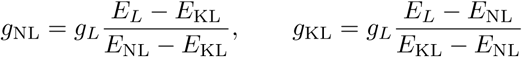

We calibrate the parameters that govern the DAG-dependent suppression of K2P channels using published experimental data from [33]. In one such experiment, DGK was over-expressed and a membrane-permeable DAG analog was applied. Once the application of this DAG analog stopped, we assume that intracellular DAG levels quickly return to baseline due to DGK over-expression, allowing K2P channels to recover to their normal open state. This assumption allowed us to compute *τ_K_* from the reported recovery time-course of the K2P conductance (Fig. 2a, left). Parameters *β_K_, K_min_* were then chosen using another experiment from the same study in which the same membrane-permeable DAG analogs were applied in a range of concentrations and the resulting efficacy of the K2P channels was measured.

**Fig. 2.**
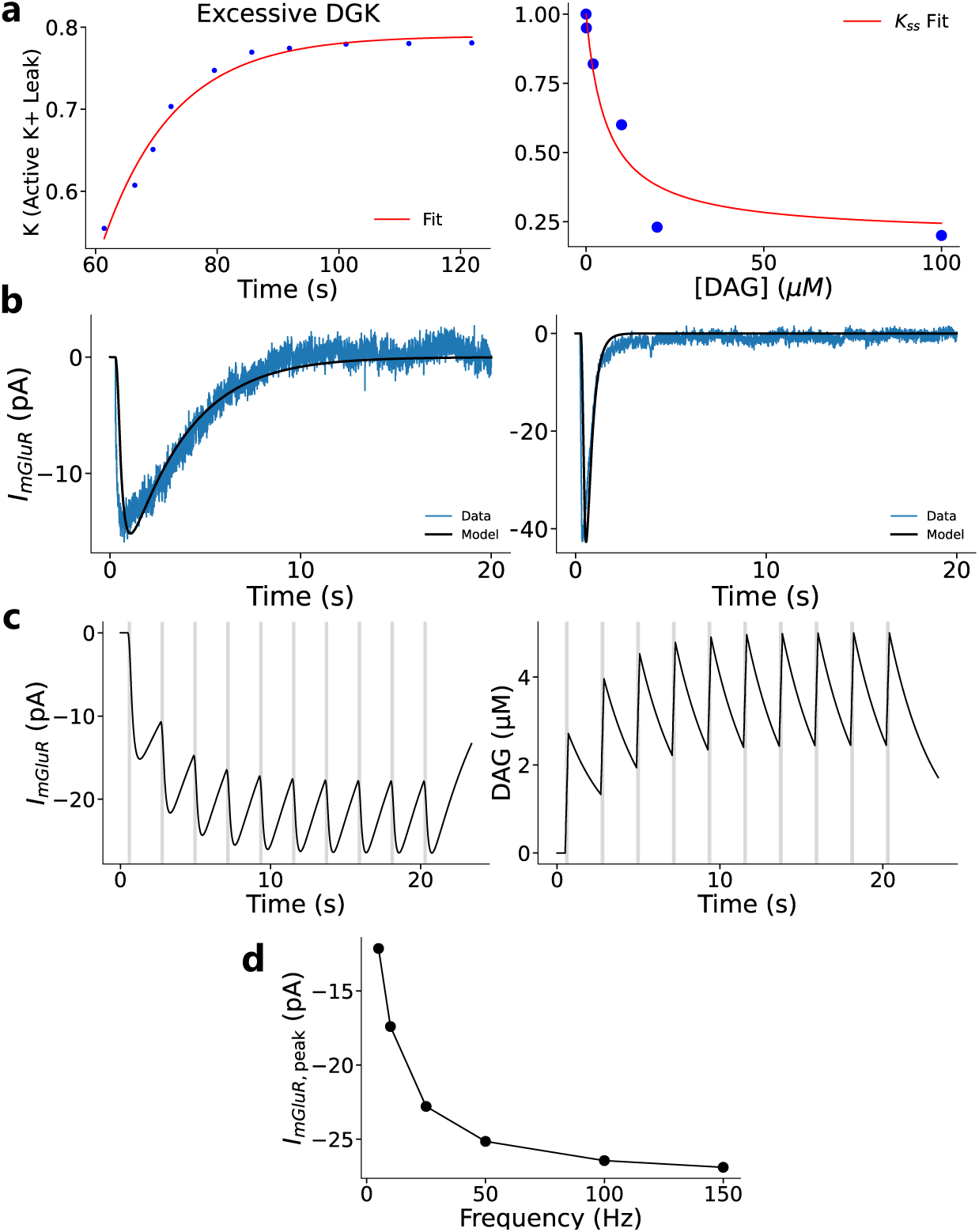
Phenomenological Model of the mGluR EPSC. **(a.)** Two experiments from [33] constrain our model parameters. Time course of K2P channel efficacy at room temperature (21°C) under excessive DGK condition (left). We assume that intracellular DAG rapidly returns to baseline, and fit a single-exponential to approximate *τ_K_* . A temperature correction (*Q*_10_ = 10) is applied to extrapolate *τ_K_* to body temperature (36.5°C), yielding *τ_K_* = 331.6 ms. Dose-response relationship between DAG concentration and K2P efficacy (right). We find *K*_min_ = 0.2 and use the steady-state analytical expression *K*_ss_ to fit *β_K_* = 5.22 × 10*^−^*^4^ *µ*M*^−^*^1^ms*^−^*^1^ (see Methods). **(b.)** mGluR EPSC recordings from cerebellar unipolar brush cells evoked by mossy fiber stimulation (20×100 Hz) [34], together with a model unipolar brush cell, fit the remaining parameters *α*_DAG_ and *τ*_DAG_. The model reproduced slow and fast EPSC waveforms with *α*_DAG_ = 0.135 *µ*M, *τ*_DAG_ = 2800 ms (left) and *α*_DAG_ = 1.6 *µ*M, *τ*_DAG_ = 130 ms (right). **(c.)** Using *α*_DAG_ = 0.135 *µ*M and *τ*_DAG_ = 2800 ms, mGluR EPSCs summate across repeated trials of presynaptic input (20 × 100 Hz, 2 s ITI, gray bar) until the peak depolarization saturates (left) as a result of DAG accumulation (right). **(d.)** Using parameters from **c**, 20 pulses of input are given and the peak mGluR EPSC is recorded as a function of the input frequency.

By fitting this data to the steady-state expression for *K* as a function of DAG concentration (Fig. 2a, right),

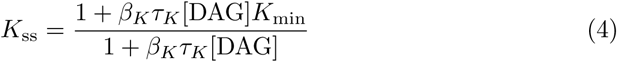

(see methods for details), the values for *β_K_* and *K*_min_ were fixed.

To account for the variability in mGluR-mediated EPSCs due to heterogeneity in DGK and PLC*β* expression in single neurons [34], we treated DAG’s decay constant (*τ*_DAG_) and the DAG production amplitude (*α*_DAG_) as free parameters. These free parameters were estimated by fitting a leaky-integrator model equipped with group I mGluR dynamics to EPSCs recorded from cerebellar unipolar brush cells in [34]. We found that our model was able to accurately reproduce the diverse set of waveforms of experimentally recorded mGluR EPSCs with these two free parameters (Fig. 2b; see Fig. S1 for additional examples), demonstrating the validity of our approach.

We next characterized the functional implications associated with the slow kinetics of mGluRs. We found that mGluR’s slow kinetics endow modulator pathways with the ability to integrate depolarizing inputs separated by several seconds. As DAG accumulates, this integration saturates (Fig. 2c). The peak magnitude of the resulting current depends on the frequency of presynaptic input (Fig. 2d), even in the presence of short-term synaptic facilitation (Fig. S2). Such slow kinetics position mGluRs as a synaptic mechanism that integrates information across seconds-long behavioral trials, a property that their ionotropic counterparts cannot realize because NMDA receptors, the slowest ionotropic receptors, generate EPSCs that decay in ∼100 ms [35].

### 2.2 Thalamocortical group I mGluRs enhance

#### somato-dendritic coupling and dendritic plateau potentials

Equipped with our phenomenological model of mGluR-mediated EPSCs, we next examine the influence of mGluR in each modulatory pathway on the trans-thalamic circuit. We begin with TC modulation, where previous experiments have shown that HOs thalamic activation of cortical group I mGluRs increases somato-dendritic coupling in L5 PNs in the sensory cortex [36], although its biophysical mechanism remains unclear. We hypothesize that this phenomenon reflects mGluR-mediated suppression of K2P channels. In cable theory of passive dendrites, a given signal attenuates along the dendro-somatic axis exponentially as exp (−*x/λ*) where *λ* is the spatial l_√_e<u>ngt</u>h constant over which a signal attenuates by a factor of 1*/e* [37]. Crucially, *λ* ∝ *R_m_*, so suppression of voltage-independent K2P channels increases membrane resistance *R_m_* and therefore *λ*. This longer space constant makes dendritic compartments more electrotonically compact and strengthens somato-dendritic coupling. Indeed, previous modeling work has demonstrated that reducing K^+^ leak channel conductance enhances dendritic integration by increasing dendritic membrane resistance [38].

To capture the effects of K2P suppression on somato-dendritic coupling, we employed a fully reconstructed multicompartmental model of a layer 5 PN from V1 that has been used extensively in previous studies [25, 26]. Guided by our phenomenological model of mGluR signaling, we varied K2P conductance in the apical dendritic compartments and quantified the resulting changes in electrotonic distance (measured in units of *λ*) along a defined somato-dendritic path. This analysis revealed that reducing K2P conductance shortens the electrotonic separation between the soma and distal apical dendrites, indicating enhanced somato-dendritic coupling (Fig. 3a). To assess how this effect extends throughout the dendrite, we additionally mapped the neuron into electrotonic space by rescaling the length of each compartment by its local space constant *λ*. This transformation provides a global visualization of dendritic coupling and shows that K2P suppression compresses electrotonic distances throughout the apical dendritic tree (Fig. 3b). To verify that these changes were also functionally relevant for arbitrary transient signals, such as those generated via synaptic and action potentials, we performed an additional morpho-electrotonic transform and generated an attenogram [39]. In this representation, distances are expressed in units of voltage attenuation such that one unit corresponds to an *e*-fold decay in voltage between any two compartments in the neuron. Consistent with our calculation of electrotonic length, decreasing *K* shortens the effective electrotonic distances in both the centripetal direction (dendrite to soma) and the centrifugal direction (soma to dendrite) (Fig. S3), indicating that suppression of K2P channel conductance by group I mGluR signaling both strengthens dendritic integration and facilitates the propagation of somatic depolarizations into the apical dendrites.

**Fig. 3.**
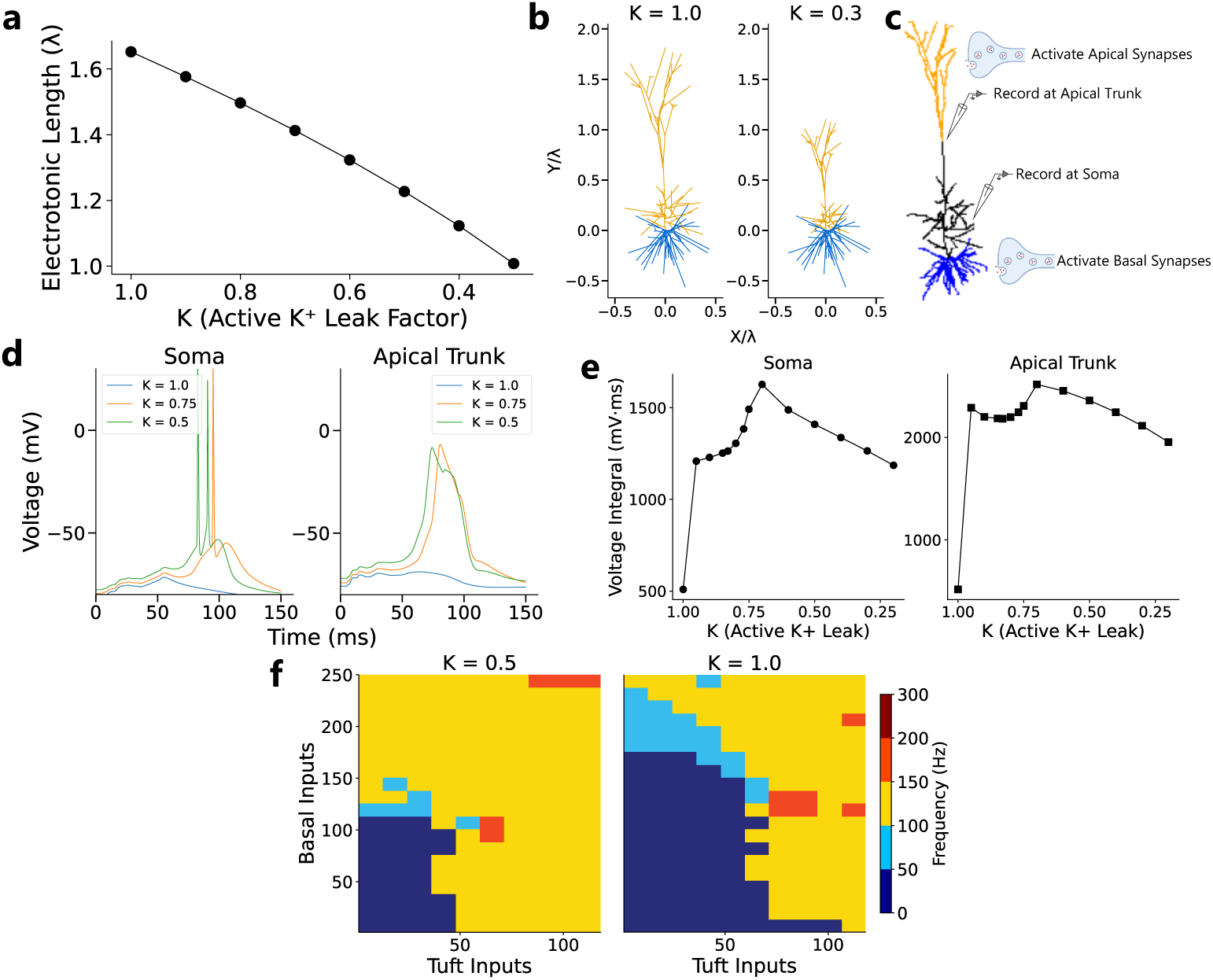
Group I mGluR-mediated K2P Suppression Modulates Activity of Dendrites in Pyramidal Neurons. **(a.)** In a fully reconstructed multicompartmental model of a L5 PN in V1, we computed the electrotonic length along a fixed path from the soma to a distal apical dendritic compartment. Reducing the K2P scaling factor *K* decreased the electrotonic length of this path, indicating enhanced somato-dendritic coupling. **(b.)** The dendritic tree is mapped into electrotonic space, where one unit of distance is defined by the space constant *λ*. Stronger K2P suppression renders the dendrite more electrotonically compact (left to right). **(c.)** Experimental paradigm probing the interaction between K2P suppression and active dendritic currents. Coincident synaptic input is given to basal dendrites and distal apical tufts while simultaneously recording from somatic and apical trunk compartments. **(d.)** With sufficiently strong mGluR-mediated K2P suppression, a fixed synaptic input (50 tuft and 50 basal inputs) is capable of triggering a dendritic Ca^2+^ spike in the apical trunk. **(e.)** Somatic and apical trunk voltage integrals as a function of decreasing *K*for fixed synaptic input (50 tuft and basal inputs). Voltage integrals changed non-linearly as *K* decreased, revealing a sharp increase in the dendritic voltage integral with *K <* 1.0 coinciding with the onset of a dendritic Ca^2+^ plateau potential. **(f.)** Greater mGluR-mediated modulation of K2Ps (*K* = 0.5) led to an increase in somatic spiking for fixed synaptic input compared to the baseline case (*K* = 1) due to the interplay between strengthened somato-dendritic coupling and dendritic plateau potentials.

Such changes in electrotonic structure through group I mGluR activation increase the gain of the neuronal input/output function to boost somatic firing, through two mechanisms that operate in tandem. First, mGluR-mediated suppression of K^+^ channels locally depolarizes the apical dendrites. Second, the accompanying enhancement of somato-dendritic coupling allows somatic depolarizations to invade the distal apical dendrites more effectively. Together, these effects reduce the synaptic input required to activate voltage-dependent currents to generate a dendritic plateau potential. With strong somato-dendritic coupling, these dendritic plateau events depolarize the soma to elicit a somatic spike that again back-propagates effectively to the dendritic compartments, establishing a ‘ping-pong’ interplay between somatic and dendritic membrane potentials that increases neuronal gain [40]. To illustrate these effects, we delivered coincident glutamatergic synaptic input to the distal apical dendrites and the soma of the multicompartmental model (Fig. 3c). Recording from compartments in the main apical trunk, we found that sufficient K2P suppression enabled a fixed pattern of coincident synaptic input to be markedly more effective in engaging voltage-gated L-type Ca^2+^ channels to facilitate a dendritic Ca^2+^ plateau (Fig. 3d, 3e). Similarly, when we applied another fixed pattern of coincident somatic and apical dendritic input and recorded from synaptically activated tuft compartments, we found that K2P suppression helped overcome the Mg^2+^ block of NMDA receptors to facilitate NMDA plateaus (Fig. S4). These dendritic plateaus, together with increased somato-dendritic coupling, synergistically increase neuronal gain and enhance somatic firing (Fig. 3f), consistent with previous experimental work [41, 42] and the established role of the HOs thalamus in selectively amplifying cortical responses to behaviorally relevant stimuli [43, 44]. This elevated firing is especially significant for L5 CT driver synapses, which are both strongly depressing and ultra-sparse [45, 46]. As such, reliably driving HOs thalamic responses *in vivo* from L5 input requires either closely spaced presynaptic events (e.g. high-firing rate burst events) rather than single isolated L5 spikes, or coincident inputs from multiple presynaptic L5 neurons to overcome the connection’s typically depressed state [47, 48]. However, because L5 CT driver synapses are ultra-sparse, short-term synaptic depression can be more easily overcome through presynaptic burst firing; indeed, we verify this prediction in a model driver synapse between a L5 PN and a HOs thalamic neuron (Fig. S5).

### 2.3 Thalamocortical mGluRs enhance perceptual detection

A large body of experimental work has shown that conscious perception in stimulus detection depends on the HOs thalamus [5, 36, 49], and stimulus detectability is correlated with the presence of dendritic plateau potentials in sensory areas [50, 51]. Given that we show that such phenomena can be regulated by group I mGluRs along TC connections, we asked whether the inclusion of mGluRs in a network of spiking neurons applied to a detection task could also shape perception.

To test this possibility, we implement a trans-thalamic network model of spiking neurons (see Methods for full details). The network is made up of two cortical areas (a lower-order sensory area and a higher-order decision area) connected directly through cortico-cortical projections and indirectly through a HOs thalamic nucleus via transthalamic pathways (see Fig. 1a for the network connectivity). In the network, the driver and modulatory synapses are distinguished by their short-term synaptic plasticity and postsynaptic receptors: driver synapses exhibit short-term depression and engage only postsynaptic ionotropic receptors, while modulatory synapses exhibit short-term facilitation and engage group I mGluRs. Beyond this connectivity, we incorporate three additional design principles in the network based on experimental work. First, recurrent excitation within the sensory area and feedback from the HOs thalamus have synaptic weights that are tuned so that, in the absence of thalamic feedback, stimulus-driven cortical activity is substantially reduced [52], making the HOs thalamus a necessary amplifier of sensory input (Fig. S6a). Second, we include a cortical inhibitory subcircuit (see Fig. 1a inset) designed so that in addition to targeting the apical dendrites of L5 PNs, HOs thalamic inputs also preferentially recruit VIP interneurons while largely ignoring SST interneurons [53], producing periods of dendritic disinhibition that are temporally aligned with TC modulatory input to enable the onset of dendritic plateau potentials (Fig. S6b) [54]. Finally, cortico-cortical weights are kept relatively weak, so the inclusion of trans-thalamic connections increases effective connectivity between the two cortical areas [3] (Fig. S6c).

We then had the network perform a stimulus detection task to investigate the behavioral consequences of TC group I mGluR activation. To simulate this task, we present a brief sensory stimulus of variable strength, given as input directly to sensory neurons for 150 ms after a fixation period of 500 ms (Fig. 4a, left). In this task, the performance is quantified by a psychometric function, namely the probability of detection as a function of input strength (e.g. contrast of a visual stimulus). Experimental work on this task has observed that the conscious detection of a stimulus is often correlated with higher-order areas undergoing an all-or-none ‘ignition’ of sustained activity, [27–29]. As such, if our network displays such sustained activity in the decision-area, the trial is classified as a hit; otherwise it is a miss (Fig. 4a, right).

**Fig. 4.**
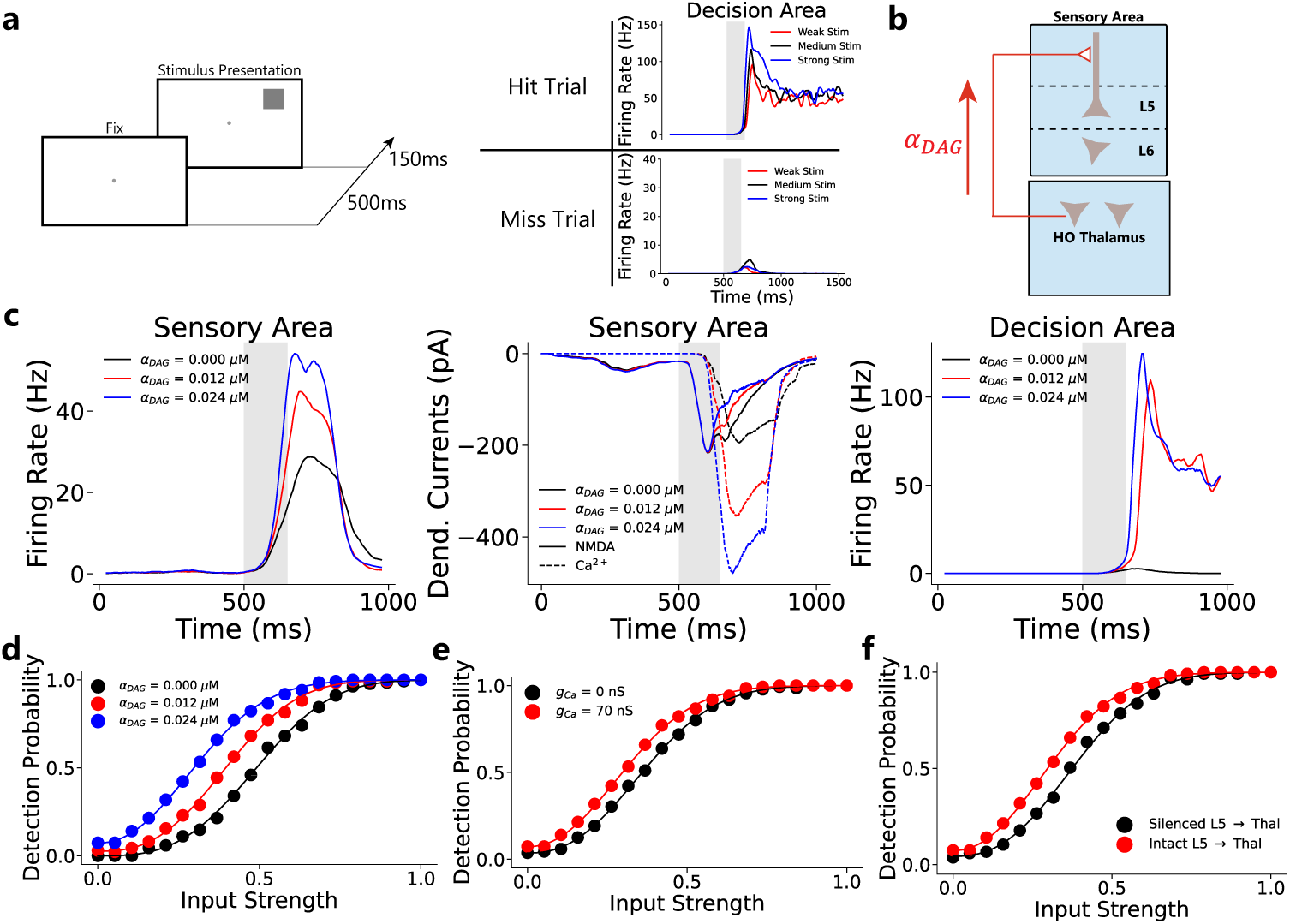
A Trans-thalamic Network of Spiking Neurons to Investigate the Role of TC mGluR Activation. **(a.)** Task Design for the Spiking Network (left): The network is presented a stimulus input for 150 ms given in the form of Poisson rates in time that obey a Gaussian Distribution after a 500 ms fixation period. Example trials at various stimulus strengths are shown (right) with stimulus presentation denoted with the gray vertical bar. In the task, a hit occurs when the decision area displays sustained activity after stimulus presentation, defined as a firing rate exceeding 20 Hz for at least 200 ms. Otherwise, the trial is a miss. **(b.)** Schematic depicting how the role of TC modulation is probed. The parameter *α*_DAG_ is increased to mimic a stronger mGluR activation resulting from thalamic activity. **(c.)** An example trial where a medium strength stimulus is presented to demonstrate the role of TC modulation. A stronger engagement of TC group I mGluRs increases the gain of sensory neurons (left) and is reflected by increases of voltage-dependent Ca^2+^ (L-type) currents (middle). This stronger input enables the higher-order decision area to undergo a dynamical bifurcation where its cortical activity is sustained (right). **(d.)** By varying across different stimulus strengths and the strength of TC modulation, we find that an increase in mGluR strength decreases the perceptual threshold. **(e.)** To confirm a synergistic interaction between mGluR and L-type Ca^2+^ channels we lesion Ca^2+^ channels and find that even with TC mGluRs (*α*_DAG_ = 0.024 *µ*M), perceptual threshold increases, aligning with results from [50, 51]. **(f.)** We also confirm the importance of L5 driver inputs to HOs thalamus in perceptual detection as with their silencing (at *α*_DAG_ = 0.024 *µ*M), the perceptual threshold is increased, aligning with results from [5].

With this setup, we next varied the strength of TC mGluR-mediated modulation to study its potential role in perception. In our network, increasing the strength of TC mGluR modulation, parameterized by the amplitude of DAG production per spike *α*_DAG_ (Fig. 4b), increased the average firing rate of L5 PNs in the sensory area, enabling stronger feedforward input that helped transition the decision area from transient to sustained high-firing activity. Consistent with our single-neuron model, this increased firing rate reflected an enhancement of dendritic Ca^2+^ plateaus (Fig. 4c). In summary, increasing the strength of mGluR modulation reduced the perceptual threshold as reflected in the leftward shift of the psychometric function (Fig. 4d).

We note that this improved detection performance relies on two synergistic effects with TC group I mGluR activation, which are crucially in alignment with previous experimental work. The first is the interaction between group I mGluR EPSCs and dendritic Ca^2+^ currents. To test whether dendritic Ca^2+^ plateau potentials were causally important in enhancing perceptual detection, we blocked Ca^2+^ currents in cortical dendrites, which increased the perceptual threshold even with intact TC mGluR modulation (Fig. 4e), in line with previous experimental work implicating dendritic Ca^2+^ signals in stimulus detection [50, 51]. The second is the contribution of feedforward trans-thalamic projections originating from L5 neurons in the sensory cortex, which not only increases the effective connectivity between the two cortical regions but also drives HOs thalamic activity to both increase its modulation of cortical gain (Fig. S7, left) and its ability to relay input to decision areas (Fig. S7, right). Silencing this feedforward thalamic driver during stimulus presentation produced perceptual deficits even when TC mGluRs were present (Fig. 4f), mirroring the experimental deficits reported by [5] and highlighting the importance of the HOs thalamus in perception.

### 2.4 Corticothalamic mGluR Shifts Thalamic Neurons from Burst to Tonic Firing Across Trials

What, then, is the role of the second modulator pathway: the projection from L6 of the sensory cortex to the HOs thalamus? This connection also strongly engages mGluRs [9], and has a key feature in which its axons collateralize with the thalamic reticular nucleus (TRN), forming a disynaptic inhibitory motif alongside the monosynaptic L6 CT modulatory connection [55]. Together with TRN’s high background firing rate [56], this motif keeps thalamic neurons hyperpolarized, de-inactivating T-type Ca^2+^ channels to favor burst firing. In such a scenario, activation of fast ionotropic receptors from L5 and L6 CT input would activate T-type Ca^2+^ channels to enable a low-threshold Ca^2+^ spike that produces a brief burst of action potentials riding its crest [57]. Switching out of this burst mode, however, requires slower depolarization sustained for at least ~100 ms, a duration that often outlasts ionotropic glutamatergic EPSCs but falls well within the seconds-long time course of mGluR-mediated depolarization [58]. Such sustained depolarization is also unlikely to arise from circuit properties alone: maintaining it through input from fast neurotransmission would require recurrent excitation among thalamic neurons, connectivity the thalamus conspicuously lacks [46]. Group I mGluRs are therefore uniquely positioned to mediate the transition between burst and tonic firing in thalamic neurons. Indeed, *in vitro* work in first-order nuclei such as LGN shows that L6 CT input can overcome TRN inhibition by engaging group I mGluRs, depolarizing relay neurons and shifting them to tonic firing [22, 59].

While this mechanism has not yet been explicitly demonstrated in the HOs thalamus, we expect the HOs thalamus to depend greatly on mGluRs, as HO thalamic neurons burst more than first-order thalamic neurons in awake behaving animals because the HO thalamus receives additional subcortical GABAergic input [60]. This increased inhibition places a greater demand on slow, summating depolarizations that mGluRs can provide to transition the HOs thalamus to tonic firing. Critically, because this depolarization decays across seconds, its summation need not be confined to a single trial of CT mGluR activation; it can accumulate across successive behavioral trials, so that the thalamic firing mode reflects the recent trial history of CT modulation.

To illustrate the role of mGluRs in mediating this transition, we model modulator synapses between L6 CT neurons and HOs thalamic neurons, represented as integrate-and-fire-or-burst (IFB) units that possess low-threshold T-type Ca^2+^ channel dynamics [57], along with the disynaptic inhibitory motif through the thalamic reticular nucleus (TRN). With these model synapses, we then simulated repeated trials of L6 CT input at 10 Hz, a level of activity observed in L6 CT neurons responding to their preferred visual stimulus [61] and one that is itself sufficient to engage group I mGluR signaling [22, 62], with a 2-second inter-trial interval (ITI) (Fig. 5a). Postsynaptic mGluR EPSCs accumulated across trials in response to L6 CT input, progressively inactivating T-type Ca^2+^ channels and transitioning thalamic neurons from burst to tonic firing modes (Fig. 5b). Notably, ionotropic NMDA currents in L6 CT connections decay with time constants of ~100 ms — far too brief to persist across the seconds-long ITI (Fig. 5b, middle), leaving mGluRs as the key mediator of cross-trial changes in the thalamic firing mode in the model synapse.

**Fig. 5.**
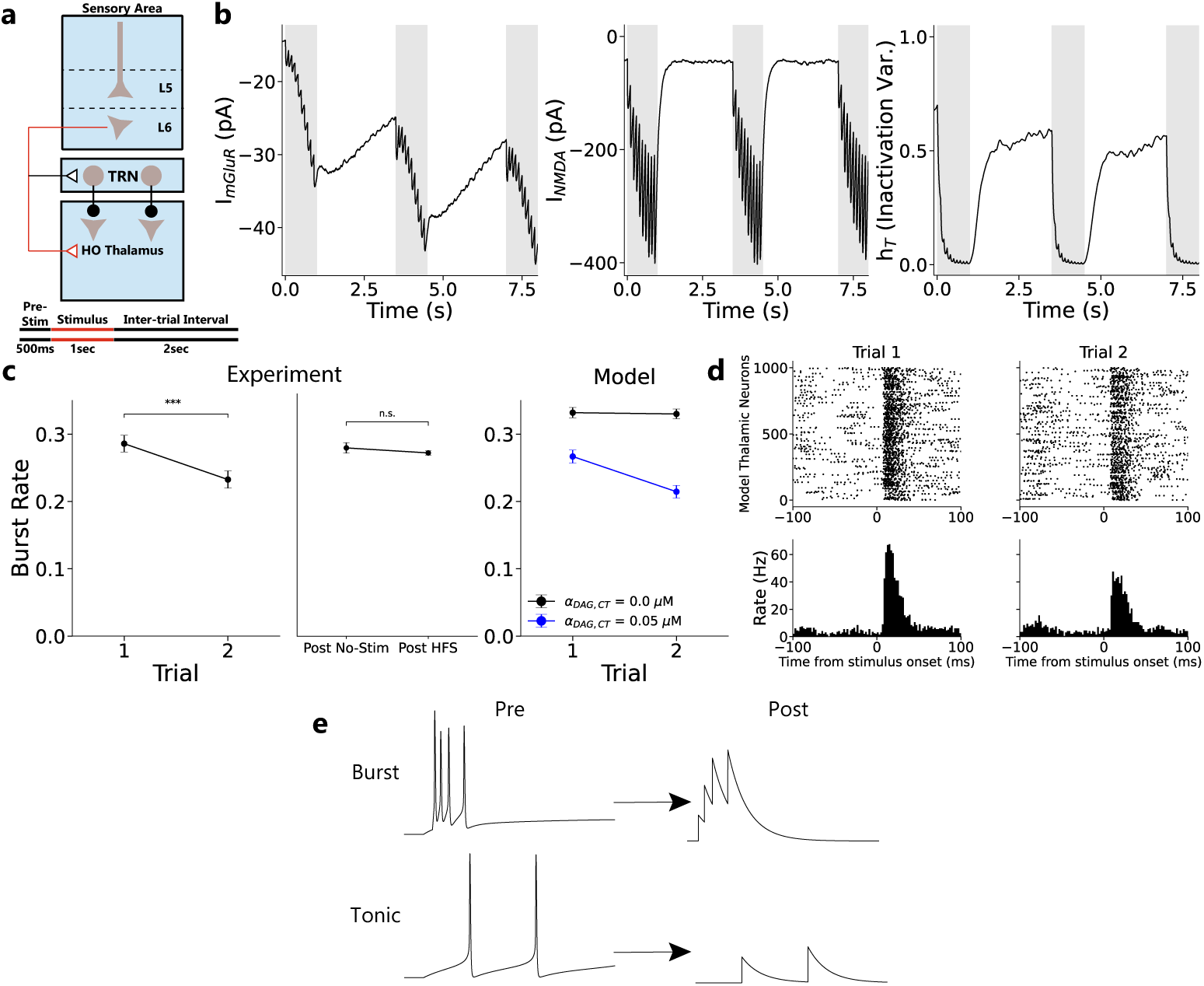
mGluR-mediated modulation of thalamic bursting across trials of cortical input. **(a.)** Simplified circuit diagram of CT modulatory synapses (top). Layer 6 CT neurons form modulatory synapses onto HOs thalamic neurons. Simultaneously, to keep thalamic neurons hyperpolarized enough to engage low-threshold T-type Ca^2+^ channels, cortical L6 neurons also make connections with the TRN, to strongly inhibit thalamic neurons. In our example simulation, L6 CT neurons fire at ∼ 10 Hz across three consecutive 1-second trials of L6 CT stimulation. Each trial begins with a 500 ms pre-stimulus period and is separated by a 2-second ITI to align with the experimental setup from [61] (bottom). **(b.)** As a result of L6 stimulation (gray bar), mGluR-mediated EPSCs accumulates across trials (left) while NMDA currents decay during the ITI due to their shorter time constant (*τ*_NMDA_ = 100 ms, middle). The depolarizing effect of group I mGluR EPSCs progressively decreases the inactivation variable *h_T_*, reducing available T-type Ca^2+^ current (right). **(c.)** Burst rate in pulvinar neurons decreases across repeated trials of L6 CT photostimulation in reanalysis of experimental data from [61] (*n* = 383; *p <* 0.001; Wilcoxon signed rank test). This decrease in thalamic bursting does not result from activity-dependent changes to synaptic strength as burst rates show no significant difference following trials with no photo-stimulation and trials where high-frequency photo-stimulation is given to L6 CT neurons (*p >* 0.2; Wilcoxon rank sum test). The observed decrease in burst firing across trials is qualitatively captured by our model synapses (right), where results are averaged across 15 simulated sequences of consecutive trials in 1000 thalamic neurons. This effect emerges only when mGluR-mediated modulation is present (*α*_DAG, CT_ = 0.05 *µ*M, blue) and is absent without it (*α*_DAG, CT_ = 0.0 *µ*M, black). **(d.)** An example sequence of trial responses of model thalamic neurons following L6 CT stimulation displays decrease in thalamic bursting at slow timescales in a raster plot of 1000 neurons (top) and their PSTH (2 ms bins, bottom) **(e.)** Functional consequence of facilitating synapses: presynaptic bursting enhances postsynaptic responses compared to tonic firing as facilitating variables do not decay as much during the inter-spike interval.

Our model results suggest a testable prediction: repeated activation of L6 CT axons across seconds-long behavioral trials should cause a observable decrease in HOs thalamic burst firing. To assess this possibility in awake-behaving animals, we reanalyzed *in vivo* extracellular recordings from the pulvinar (a HO visual thalamic nucleus) in awake-behaving mice following optogenetic photostimulation of L6 CT neurons in V1 [61]. We examined the firing patterns of pulvinar neurons across trials of 10 Hz L6 photo-stimulation, separated by a 1.5 to 2 second ITI. In our analysis, each analyzed sequence of trials was preceded by a trial without optogenetic input to allow any slow depolarizing processes to decay. Consistent with mGluR-mediated EPSCs shifting thalamic neurons toward tonic firing at slow timescales, we observed a significant decrease in the rate of thalamic bursts in consecutive behavioral trials (*n* = 383; *p <* 0.001; Wilcoxon signed rank test) (Fig. 5c, left). Although we cannot definitively attribute this observed decrease to group I mGluR activation, alternative mechanisms are unable to explain this result. A key scenario that we consider is activity-dependent synaptic plasticity that persists over seconds, including short-term facilitation in L6 CT connections, short-term depression of disynaptic TRN inhibition [3, 63, 64], synaptic augmentation, or post-tetanic potentiation [65], which could potentially increase depolarizing effects in a subsequent trial. However, the rate of thalamic bursts during 10 Hz photo-stimulation trials preceded by high-frequency L6 stimulation (20/40 Hz) (which would strongly engage these plasticity mechanisms) did not differ significantly from trials preceded by no optogenetic stimulation (*p >* 0.2; Wilcoxon rank-sum test) (Fig. 5c, middle), suggesting that short-term plasticity at these synapses does not persist meaningfully across behavioral trials *in vivo*. By contrast, subjecting our model synapses equipped with postsynaptic group I mGluRs to an identical stimulation protocol reproduced the observed decrease in thalamic burst firing across trials of L6 CT stimulation, and this effect was abolished by selectively blocking group I mGluRs (Fig. 5c, right). Model raster plots and peristimulus time histograms (PSTHs) further illustrate the reduction of thalamic burst firing across consecutive trials (Fig. 5d). Together, our results identify mGluR-mediated integration as a parsimonious explanation for the observed decrease in thalamic bursting across trials.

What is the relevance of this transition in thalamic firing modes in perception? We speculate that the ability to control transitions between burst and tonic firing can provide an effective mechanism for regulating information flow in trans-thalamic circuitry in a history-dependent manner, particularly given the strongly facilitating nature of TC feedback connections onto sensory cortices. In facilitating synapses, closely spaced events such as thalamic bursts enhance postsynaptic activation of glutamate receptors, because facilitation variables do not decay nearly as much across short inter-spike intervals (Fig. 5e). Thus, thalamic burst firing could enhance HOs thalamic modulation of cortical PNs. However, as mGluR EPSCs accumulate from repeated CT activation across behavioral trials, thalamic neurons shift from burst to tonic firing, reducing the strength of subsequent TC modulation. Importantly, although bursting strengthens TC feedback, we note that in our model, thalamic bursting is not required for TC mGluR activation and its ability to enhance the feedforward propagation of sensory information. In particular, we found that increasing the strength of TC mGluR modulation increased the firing of sensory neurons (Fig. S8a) to enhance conscious detection even when thalamic neurons fired tonically through a constant 30 pA depolarizing current injection (Fig. S8b).

### 2.5 Corticothalamic mGluRs implement a ‘thalamic wake-up call’

Given the interplay between TC and CT modulation on single-neurons, we propose that modulatory connections in the trans-thalamic circuit work in concert with each other to implement behaviors reminiscent of the long-standing thalamic wake-up call hypothesis. In this scenario, thalamic bursts enhance cortical firing in response to novel and salient stimuli, thus promoting their detection [58, 60, 66]. In our model, we propose that this “thalamic wake-up call” is implicitly realized in the trans-thalamic circuit, where an initial burst of activity in the HOs thalamus more effectively engages activation of postsynaptic TC receptors (including mGluRs) in sensory neurons to enhance their gain. In doing so, this strengthens feedforward input to neurons in the decision circuit to enable an all-or-none ‘ignition’ of sustained activity, a dynamical motif indicative of conscious perceptual detection. However, as the stimulus is presented more frequently and its novelty decreases, repeated activation of the same CT synapses accumulates mGluR EPSCs to shift HOs thalamic neurons into a tonic firing mode, reducing the HOs thalamus’s ability to overcome the facilitating nature of its feedback to early sensory areas. Consequently, thalamic enhancement of dendritic plateaus progressively weakens, consistent with a diminished need for a thalamic “wake-up” signal due to the decreased novelty of the sensory stimulus.

Motivated by the slow kinetics of mGluR-mediated EPSCs and their role in regulating thalamic firing modes, we utilize our developed network model and repeatedly present a stimulus of identical strength to the network across sequential trials (2 s ITI, Fig. 6a). Despite the stimulus strength remaining unchanged, the mean firing rate of L5 neurons in the sensory cortical area progressively decreased across behavioral trials, reducing the mean firing rate in the downstream decision area following stimulus presentation (Fig. 6b). As predicted, we found an accumulation of CT mGluR EPSCs, whose slow depolarization progressively inactivated T-type Ca^2+^ channels in HOs thalamic neurons, biasing them away from burst firing and toward a tonic firing regime. In this tonic firing regime, TC modulation weakened and the recruitment of voltage-dependent NMDA and L-type Ca^2+^ currents was reduced in the apical dendrites of sensory pyramidal neurons, thus decreasing the gain of these neurons (Fig. 6c). At the behavioral level, these circuit-level adaptations manifested as a progressive decline in detection performance across repeated stimulus presentations, with perceptual thresholds increasing across behavioral trials (Fig. 6d). We then reasoned that if this adaptation is caused by depolarization from CT mGluR EPSCs, then lengthening the ITI and allowing the mGluR-mediated EPSC to decay between stimuli should lessen the observed cross-trial adaptation of perceptual detection. Indeed, we found that extending the ITI from 2 s to 5 s allowed substantial decay of the CT mGluR EPSC between trials (Fig. 6f, inset) and the adaptive rightward shift of psychometric function was nearly abolished, with the detection performance of Trial 2 closely overlapping Trial 1 (Fig. 6e). Examining a wider range of ITIs, we found that the perceptual detection threshold (50% hit rate) of the 2nd sequential trial decreased as the

**Fig. 6.**
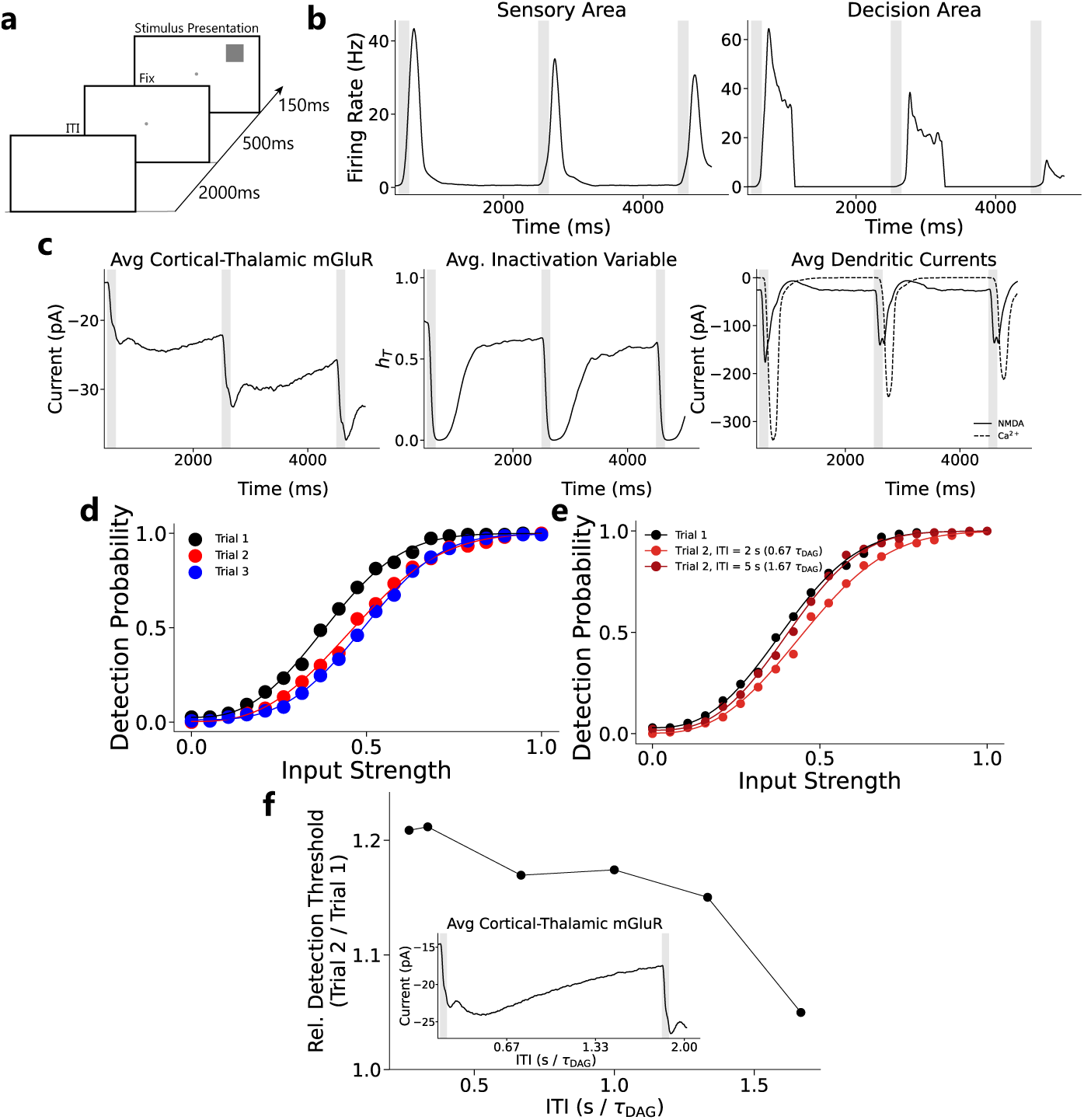
CT mGluR activation mediates cross-trial adaptation. **(a.)** To study CT mGluR’s effects at seconds-long timescale, we introduce a 2 second inter-trial interval (ITI), between trials in our detection task and fix TC mGluR strength to *α*_DAG_ = 0.012 *µ*M. **(b.)** Trial-averaged firing rates (15 simulated sequences of three consecutive trials) in the sensory area (left) and decision area (right) in response to inputs of medium strength. Gray vertical bars mark stimulus presentations, with input strength drawn from a Gaussian of fixed mean. Sensory firing decreases across trials, in turn diminishing the post-stimulus response in the decision area. **(c.)** Trial-averaged traces from the same sequences: CT mGluR EPSCs accumulate across trials (left), reducing the T-type Ca^2+^ inactivation variable and shifting thalamic neurons from burst to tonic firing (middle), which lowers average dendritic currents in sensory cortex (right). **(d.)** With CT mGluR activation (*α*_DAG,CT_ = 0.05 *µ*M), the psychometric function shifts progressively rightward across trials, indicating an increasing perceptual threshold. **(e.)** Elongating the ITI from 2 seconds to 5 seconds nearly abolishes the cross-trial shift in the psychometric function. **(f.)** Across various seconds-long ITIs, we find that the Trial 2 perceptual threshold (the input strength with 50% detection rate) decreases with ITI length. Inset: This change in detection threshold is mediated by the decay of mGluR-mediated EPSCs during a longer, 5 s, ITI as the slowest dynamic, *τ*_DAG_, is only 3 s.

ITI lengthened (Fig. 6f), highlighting how the temporal statistics of incoming stimuli help set the degree to which CT modulation attenuates detection according to the magnitude of the mGluR EPSC.

## 3 Discussion

In this work, we developed a modeling framework that spans three biological levels to shed light on into how the engagement of group I mGluRs shapes computation and cognition in trans-thalamic circuitry. At the molecular level, we introduced a biophysically constrained model of mGluR signaling capable of reproducing the diverse slow EPSCs resulting from mGluR-mediated suppression of K2P channels.

Building on the phenomenological kinetic model, we examined the functional consequences of group I mGluR signaling in both TC and CT modulatory connections at the single-neuron and network levels. In TC modulatory connections, we find that group I mGluR activation in distal apical dendrites enhanced somato-dendritic coupling and promoted the generation of dendritic plateau potentials in a fully reconstructed L5 PN multicompartmental model. These dendritic plateaus in sensory neurons, in turn, boosted the feedforward propagation of sensory information in a trans-thalamic network of spiking neurons. Importantly, our model connects mGluRs to phenomena that experimental work has shown to be crucial for performance in a detection task. First, driving input from cortex to thalamus has been shown to increase detection performance [5], which, in the context of our model, excites HO thalamic neurons and increases TC modulation over sensory neurons. Second, dendritic plateau potentials have previously been identified as crucial for detection [50, 51]; in our model, these plateaus are enhanced by mGluR activation.

Furthermore, in CT modulatory connections, we find that HOs thalamic neurons transition from burst to tonic firing as a result of the accumulation of mGluR-mediated EPSCs that were activated by CT modulator inputs across behavioral trials, over many seconds. This prediction was supported by a reanalysis of published *in vivo* recordings in the pulvinar from behaving animals [61]. At the network level, this shift towards tonic firing in HOs thalamic neurons reduced their modulatory influence on the sensory cortex and attenuated detection performance. By connecting levels of biological abstraction, our model shows how molecular machinery (mGluR receptors) can be causally linked through neural circuits to implement a dynamic, history-dependent regulation of cortical gain capable of shaping perception.

The present study suggests several directions for future investigation. At the single-neuron level, we propose that the absence of group I mGluR-mediated K2P suppression causes the pronounced somato-dendritic decoupling observed during general anesthesia, which dramatically reduces thalamic activity [36]. This prediction can be tested by pharmacological blockade of K2P channels to assess whether somato-dendritic coupling is restored even in the presence of group I mGluR antagonists. However, we note that group I mGluRs are known to modulate additional ion channels that can contribute to this phenomenon [18], including other K^+^ currents [67] and HCN channels [68] whose suppression has been experimentally and theoretically shown to be capable of increasing the electrotonic compactness of dendrites [69, 70]. Our focus on K2P suppression may therefore underestimate the full extent of mGluR’s ability to enhance somatodendritic coupling, and future experiments in tandem with computational modeling should parse the relative contributions of these targets.

At the network level, our demonstration that the thalamic firing mode governs the effectiveness of TC gain modulation helps address a longstanding mystery: the functional role of thalamic bursting in cognition. Although thalamic bursting has historically been associated with sleep and disconnected states, recent causal evidence demonstrates its relevance for attention and perception in awake behaving animals [71], with thalamic bursts in pulvinar being associated with enhanced detection performance, in line with our network’s prediction that bursting increases thalamus’s modulatory influence over cortex. In our model, thalamic bursting is triggered by feed-forward sensory input under conditions in which group I CT mGluR EPSCs are weak, but understanding what other excitatory/inhibitory and neuromodulatory sources can affect burst firing remains crucial to gaining a complete understanding of thalamic function in cortical processing. In addition, recent anatomical work has characterized extensive feedback projections from higher-order cortical areas and motor cortices to HOs thalamic nuclei [12], suggesting that the HOs thalamus can serve as a hub that integrates ascending sensory signals with top-down contextual information [72] and efference copies [2]. Experimental evidence has further revealed that HOs thalamic activity can encode crucial cognitive variables such as decision confidence [73] and sensory prediction errors [44]. Together, this anatomical connectivity and the encoding of these important cognitive variables suggest that the HOs thalamus is well-positioned to flexibly affect cortical processing according to task demands. Determining how these diverse inputs interact to dynamically drive and modulate thalamic firing and, in turn, shape cortical sensory representations represents an important direction for future computational and experimental investigations.

Finally, we note that our model thus far has addressed only the contributions of group I mGluRs in trans-thalamic circuitry. However, within the class of all mGluRs, there are two other major sub-groups, group II and group III mGluRs. These group II and III mGluRs can act pre- and postsynaptically to decrease neuronal excitability [14, 74] by activating G_i/o_ proteins that can engage G protein-coupled inwardly-rectifying K^+^ channels [75]. It is noteworthy that at a behavioral level, postsynaptic group II mGluR3 in the primate cortex is implicated in higher cognition and intelligence [76, 77]. Extending our present framework to incorporate these additional targets and capture the potential interplay between these diverse mGluR subtypes will be an important direction for future work.

## 4 Methods

### 4.1 Designing a Model of the Molecular Signaling in group I mGluRs

Our first goal is to design a computationally tractable phenomenological model of group I mGluR mediated EPSCs. The model is designed to capture the EPSC’s long time-course and DAG-dependent K2P suppression, which are described by two dynamical equations:

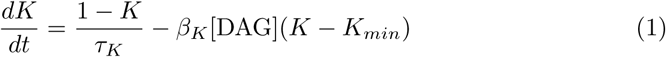

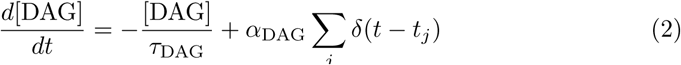

Here DAG reduces K2P channel availability to a minimum value of *K_min_*, with K2P channels recovering with time constant *τ_K_*. Pre-synaptic spikes (represented by the Kronecker delta function) increase DAG by *α*_DAG_, which decays with *τ*_DAG_.

The resulting EPSC arises from K2P suppression, assuming that the leak current is composed of non-selective cation and K^+^ components:

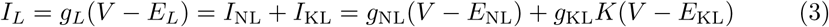

with

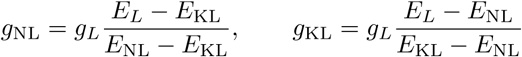

Here, *g*_NL_ and *g*_KL_ denote the non-selective cation and K^+^ leak conductances. We set the reversal potentials of the non-selective cation leak channels and the K^+^ leak channels as *E*_NL_ = 0 mV [78] and *E*_KL_ = − 100 mV [32], respectively.

The parameters governing the DAG-dependent suppression of K2P were constrained by experiments [33]. *τ_K_* was estimated from the recovery of K2P conductance after the application of a membrane-permeable DAG analog under conditions of DGK over-expression; a Q_10_ = 10 correction for K2P [79] was applied to the fitted value of *τ_K_* so that it can be used to model dynamics at body temperature (36.5*^◦^*C). We then used the steady-state dose-response curve of K2P suppression as a function of DAG concentration and found that *K_min_* = 0.2 while fitting *β_K_* to the steady-state expression of *K*:

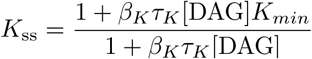

by applying the BFGS algorithm [80]. These steps ultimately give us *τ_K_* = 331.6 ms *β_K_* = 5.22 10*^−^*^4^*µ*M*^−^*^1^ms*^−^*^1^. The remaining parameters, *α*_DAG_ and *τ*_DAG_, were fit to mGluR EPSC traces in cerebellar unipolar brush cells [34], using a leaky integrator model. Mathematically, the leaky integrator this model is given by:

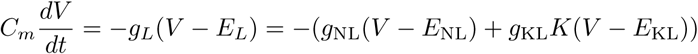

In our simulations, we used *g_L_* = 2.6 nS, *C_m_* = 16.6 pF, *E_L_* = −65 mV, values which are in line with passive properties observed in [81].

### 4.2 Multicompartmental Model

We employed a fully reconstructed multicompartmental model of a layer 5 pyramidal neuron originally introduced by [25], which faithfully reproduces key dendritic non-linearities, including Ca^2+^ plateau potentials and backpropagating action potentials (bAPs). Model parameters were taken from a subsequent study [26], where they were fit to electrophysiological recordings. This parameterization also incorporates NMDA and AMPA synapses distributed across basal and apical dendrites, with a maximum NMDA-to-AMPA conductance ratio of 1:1 per synapse.

In our simulations, the leak current was decomposed into non-selective cation and K^+^ components as described in Eq. (3), with the K^+^ leak conductance scaled by the mGluR-dependent factor *K*. After the neuron reached steady state, we delivered coincident synaptic input to clustered basal and apical dendritic compartments and recorded membrane voltage in the soma and apical trunk to capture dendritic Ca^2+^ plateau potentials. We also similarly delivered a second pattern of coincident input to the same basal/apical dendritic compartments and recorded the voltage responses in activated apical compartments to capture dendritic NMDA plateau potentials.

Then, to understand how changing *K* could change the nature of dendritic signaling in the neuron, we computed the total electrotonic length *Z* of a given path from a somatic to a distal dendritic compartment as

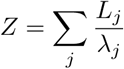

where *L_j_* is the length and *λ_j_* is the local space constant of a given compartment, *j*. In a passive cable, this is given as

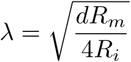

where *d* denotes the diameter of the compartment and *R_m_* and *R_i_* are the membrane and axial resistances, respectively [37]. In addition, we characterized the effect of changing *K* for the entire dendritic tree by generating graphical representations of the neuron in electrotonic space. For the classical electrotonic representation, compartment lengths *L_j_* were scaled by their local space constant, *λ_j_*. To generate attenograms that could capture the effective electrotonic changes in dendrites of arbitrary geometry, compartment lengths were instead scaled by their log-attenuation *A_ij_* [39],

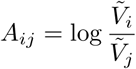

where *V*^^^*_i_* and *V*^^^*_j_* denote the voltage changes at the injection site *i* and at the recording location *j*.

### 4.3 Experimental Data Analysis

We analyzed *in vivo* extracellular recordings from pulvinar neurons in six awake, head-fixed mice from [61]. In this experimental protocol, mice were free to locomote while viewing square-wave drifting gratings or a gray screen. Each trial began with a 500 ms pre-stimulus period followed by 2 seconds of visual presentation. Optogenetic stimulation of L6 CT neurons in V1 was given 500 ms after stimulus onset and lasted 1 second at varying stimulation frequencies. The trials were separated by a 1.5 to 2 second ITI.

To identify units that were activated by L6 CT photostimulation, we constructed each unit’s PSTH of spike counts (5 ms bins) across trials of 10 Hz or 20 Hz photo-stimulation. A response threshold was defined as three standard deviations above the mean visually evoked response during the one-second window corresponding to the photostimulation period on no-light trials. Units were kept for further analysis if they met either of the following criteria: (1) at least one bin exceeded the response threshold within 50 ms of the first or second photostimulation pulse; or (2) in the absence of a significant response to either of the first two pulses, at least half of all subsequent pulses were followed by bins that exceeded the response threshold.

To quantify burst firing for all remaining units, we first classified which spiking events were part of bursts. Following convention, a burst was defined as a sequence of spikes in which the first spike was preceded by an inter-spike interval (ISI) ≥ 100 ms and followed by at least one spike with an ISI ≤ 4 ms. Subsequent spikes were considered part of the same burst if they were preceded by an ISI ≤ 4 ms. Burst rate was then computed as the ratio of burst events to the total number of spiking events. An identical calculation was done on model HOs IFB thalamic units to illustrate how CT mGluR EPSCs can reduce thalamic bursting.

To assess the potential cross-trial effects of L6 CT stimulation, we identified sequences of behavioral trials that meet several criteria. In all analyses, we restricted the trials to those during which the animal remained stationary, because locomotion and arousal are known to strongly modulate thalamic firing [82]. Because visual stimulus conditions and optogenetic frequencies had been independently randomized in the original experiment, we searched for sequences in which the same photostimulation frequency occurred in consecutive trials by chance, allowing us to evaluate cross-trial effects at a fixed stimulation strength. To minimize the residual effects of lingering slow synaptic dynamics, we further required each sequence to be preceded by a trial without optogenetic L6 CT activation or visual stimulus presentation. Although qualifying sequences were relatively infrequent, we found that consecutive two trial sequences provided us with 383 units to perform statistical analysis.

Within these sequences, we focus on 10 Hz photostimulation, as this regime most closely approximates the *in vivo* firing rates of L6 CT neurons responding to preferred visual stimuli [61], while being at a level capable of activating mGluRs [22, 62]. To further test the contribution of short-term synaptic plasticity to pulvinar firing, we additionally identified sequences in which 10 Hz trials were preceded by high-frequency stimulation (20/40 Hz) and compared these to 10 Hz trials that were preceded by a trial with no stimulation.

### 4.4 Trans-thalamic Network of Spiking Neurons

#### 4.4.1 Neurons

Our trans-thalamic network of spiking neurons is comprised of three interconnected areas: a first-order cortical area representing sensory processing (sensory area), a higher-order cortical decision area associated with decision making (decision area), and a HOs thalamic area that indirectly connects the two cortical regions. The sensory area possesses different populations of neurons that correspond to L5 and L6 neurons. Within L5, we have a population of 1600 excitatory neurons and 400 inhibitory neurons. Of the inhibitory neurons, 160 were soma-targeting PV interneurons, 160 were dendrite-targeting SST interneurons, and 80 were SST-targeting VIP interneurons. Within L6, we have a population of 1000 excitatory neurons and 200 inhibitory neurons. On the other hand, the decision area makes no distinction between neurons from different layers and uses the same set-up as L5 Neurons in the sensory areas for simplicity (1600 excitatory Neurons, 160 PV interneurons, 160 SST interneurons, 80 VIP interneurons). The HOs thalamus possesses 2000 excitatory neurons and receives GABAergic input from the TRN which is made up of 400 inhibitory neurons.

L5 excitatory neurons (and HO decision-area neurons) are modeled as two-compartment leaky-integrate-and-fire neurons (LIF) equipped with a dendritic Ca^2+^ current based on [41]. Mathematically, their dynamics are given by the following dynamical equations:

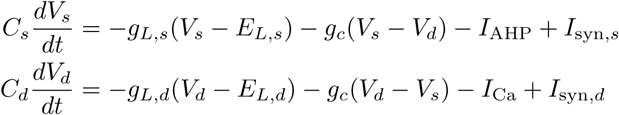

The adaptation current (AHP) [41] and the dendritic Ca^2+^ current [83] were given by:

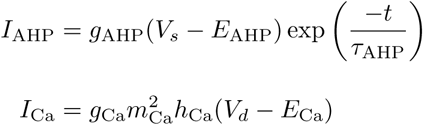

For the AHP current, *t* represents the time from the neuron’s most recent spike. The Ca^2+^ gating variables *m*_Ca_ and *h*_Ca_ satisfy first-order kinetics:

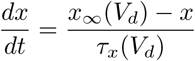

where 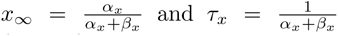 . For the activation variable *m*_Ca_, *α_m_* = 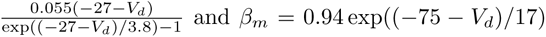. For the inactivation variable *h*_Ca_, *α_h_*= 0.000457 exp((−13 − *V_d_*)*/*50) and 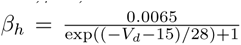 Within these equations, *C_s_, C_d_* describe the capacitance of the somatic and dendritic compartments, respectively. *E_L_, E*_Ca_*, E*_AHP_ represents the reversal potentials of the leak, Ca^2+^ (L-type), AHP currents, respectively. *g_L_, g*_Ca_*, g*_AHP_ denotes the conductance of the leak, Ca^2+^ (L-type), AHP currents, respectively. The parameter *g_c_* represents the axial conductance between the two compartments.

To capture K2P-mediated changes in somato-dendritic coupling in the reduced two-compartment model, we scaled the amplitude of the bAP to match the centrifugal propagation observed in the multicompartmental model (Fig. S9A). In this framework, a bAP is implemented as a voltage kick to the dendritic compartment triggered by a somatic spike, with its amplitude modulated by the fraction of open K2P channels, *K*. We derived the relationship between *K* and the amplitude of bAP by varying *K* in the apical compartments of the multicompartmental model, eliciting a somatic spike, and measuring the resulting change in dendritic voltage in the apical trunk. A linear fit to these measurements yielded 16.431 − 4.783*K* (Fig. S9B).

L6 excitatory neurons were modeled as single-compartment LIF neurons. Mathematically, their dynamics are given by:

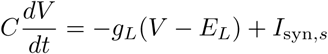

HOs thalamic neurons were modeled as single-compartment IFB neurons [57]. Their dynamics are given by:

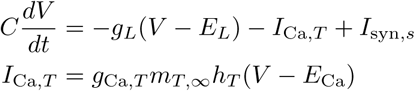

The gating variables are given by *m_T,∞_* = *H*(*V* − *V_h_*) where *H* is the Heaviside step function and *V_h_*is the threshold responsible for the activation of burst spiking. The inactivation variable *h* has its dynamics given by:

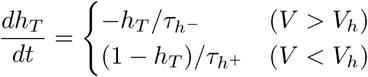

All inhibitory neurons were modeled as single-compartment LIF neurons equipped with adaptation. Mathematically, their dynamics are modeled by the following dynamical equation:

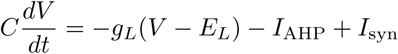

where, like excitatory neurons, the adaptation current is given by: *I*_AHP_ = *g*_AHP_(*V –* 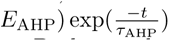, where *t* is the time following a spike event.

Background poisson input is given to AMPA synapses for each neuron so that their background firing rate ranges from 1 to 8 Hz. An exception is the TRN where the background firing rate is around 35 Hz, aligning with [56].

A complete list of parameters associated with each neuronal model and the experimental references (if present) used to constrain their values is found in Table 1.

**Table 1:** Neuronal Parameters in Trans-thalamic Network.

| Parameter | Value | Reference |
| --- | --- | --- |
| **Layer 5 PNs** |  |  |
| $C_s$ | 260 pF | [41] |
| $C_d$ | 120 pF | [41] |
| $g_{L,s}$ | 20 nS | [41] |
| $g_{L,d}$ | 23.26 nS | [41] |
| $g_c$ | 15.38 nS | [41] |
| $E_{L,s}$ | -70 mV | [41] |
| $E_{L,d}$ | -60 mV | [41] |
| $g_{Ca}$ | 70 nS | [41] |
| $E_{Ca}$ | 120 mV | [41] |
| $g_{AHP}$ | 4 nS | [41] |
| $E_{AHP}$ | -90 mV | [41] |
| $\tau_{AHP}$ | 80 ms | [41] |
| $V_{\text{threshold}}$ | -47 mV | [41] |
| $V_{\text{reset}}$ | -52 mV | [41] |
| $\tau_{\text{refractory}}$ | 1 ms | [41] |
| **Layer 6 PNs** |  |  |
| $C$ | 75.6 pF | |
| $g_L$ | 4.3 nS | |
| $E_L$ | -70 mV | |
| $V_{\text{threshold}}$ | -50mV | |
| $V_{\text{reset}}$ | -55mV | |
| $\tau_{\text{refractory}}$ | 2 ms | |
| **Thalamic Neurons** |  |  |
| $C$ | 340 pF | |
| $g_L$ | 12 nS | |
| $E_L$ | -63 mV | |
| $g_{Ca,T}$ | 15 nS | |
| $\tau_{h-}$ | 20 ms | [57] |
| $\tau_{h+}$ | 100 ms | [57] |
| $V_h$ | -60 mV | [57] |
| $E_{Ca}$ | 120 mV | |
| $V_{\text{threshold}}$ | -35 mV | [57] |
| $V_{\text{reset}}$ | - 50 mV | [57] |
| **PV Interneurons** |  |  |
| $C$ | 60 pF | |
| $g_L$ | 8.33 nS | |
| $E_L$ | -70 mV | |

Table 1 – *Continued from previous page*
| Parameter | Value | Reference |
| --- | --- | --- |
| $g_{AHP}$ | 0 nS | |
| $V_{\text{threshold}}$ | -50 mV | |
| $V_{\text{reset}}$ | -55 mV | |
| $\tau_{\text{refractory}}$ | 2 ms | |
| **SST Interneurons** |  |  |
| $C$ | 70 pF | |
| $g_L$ | 4 nS | |
| $E_L$ | -65 mV | |
| $g_{AHP}$ | 0.3 nS | |
| $\tau_{AHP}$ | 500 ms | |
| $E_{AHP}$ | -90 mV | |
| $V_{\text{threshold}}$ | -50 mV | |
| $V_{\text{reset}}$ | -55 mV | |
| $\tau_{\text{refractory}}$ | 4 ms | |
| **VIP Interneurons** |  |  |
| $C$ | 56 pF | |
| $g_L$ | 2.77 nS | |
| $E_L$ | -65 mV | |
| $g_{AHP}$ | 0.4 nS | |
| $\tau_{AHP}$ | 500 ms | |
| $E_{AHP}$ | -90 mV | |
| $V_{\text{threshold}}$ | -50 mV | |
| $V_{\text{reset}}$ | -55 mV | |
| $\tau_{\text{refractory}}$ | 4 ms | |
| **TRN Interneurons** |  |  |
| $C$ | 80 pF | |
| $g_L$ | 3.79 nS | |
| $E_L$ | -65 mV | |
| $g_{AHP}$ | 0 nS | |
| $V_{\text{threshold}}$ | -50 mV | |
| $V_{\text{reset}}$ | -55 mV | |
| $\tau_{\text{refractory}}$ | 2 ms | |

#### 4.4.2 Synapses

The connectivity of the trans-thalamic network is based on Fig. 1. Recurrent and interareal connections were kept sparse, with connection probabilities of approximately 6–11%. An exception was made for projections from L5 PNs to the HOs thalamus, which were set to be extremely sparse such that the expected number of synapses per target neuron was approximately 2.5, consistent with experimental reports [43]. See Table 2 for a complete set of connection probabilities. In our model, we include four synapse types: AMPA, NMDA, GABA, and mGluR. Following [17], AMPA and GABA are modeled as linear:

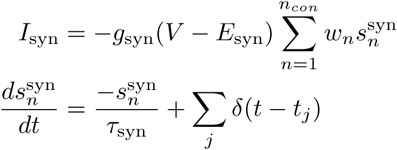

**Table 2.** Network Connectivity Matrix - Connection Probabilities

| Pre \ Post | L5 | L6 | Dec | Thal FF | Thal FB | PV | SST | VIP | PV,L6 | PV,D | SST,D | VIP,D | TRN |
| --- | --- | --- | --- | --- | --- | --- | --- | --- | --- | --- | --- | --- | --- |
| L5 | 0.11 | - | 0.0625 | 0.0015625 | 0.0015625 | 0.1 | 0.1 | - | - | 0.1 | 0.1 | - | 0.1 |
| L6 | - | 0.1 | - | 0.0625 | 0.0625 | - | - | - | 0.1 | - | - | - | 0.1 |
| Dec | - | - | 0.11 | - | - | 0.1 | 0.1 | - | - | 0.1 | 0.1 | 0.1 | - |
| Thal FF | - | - | 0.1 | - | - | - | - | - | - | - | - | - | - |
| Thal FB | 0.1 | - | - | - | - | - | - | 0.1 | - | - | - | - | - |
| PV | 0.1 | - | - | - | - | 0.1 | - | - | - | - | - | - | - |
| SST | 0.1 | - | - | - | - | 0.1 | - | - | - | - | - | - | - |
| VIP | - | - | - | - | - | - | 0.1 | - | - | - | - | - | - |
| (PV, L6) | - | 0.1 | - | - | - | - | - | - | - | - | - | - | - |
| (PV, decision) | - | - | 0.1 | - | - | - | - | - | - | 0.1 | - | - | - |
| (SST, decision) | - | - | 0.1 | - | - | - | - | - | - | 0.1 | - | - | - |
| (VIP, decision) | - | - | - | - | - | - | - | - | - | - | 0.1 | - | - |
| TRN | - | - | - | 0.1 | 0.1 | - | - | - | - | - | - | - | - |

where *s^syn^*is the gating variable at the *n*th synapse, *w_n_* is the synaptic weight at the *n*th synapse, *g_syn_* is the maximum synaptic conductance of the synaptic current, and *E_syn_* is the synaptic reversal potential.

NMDA receptors include a voltage-dependent magnesium block and their dynamics are given by:

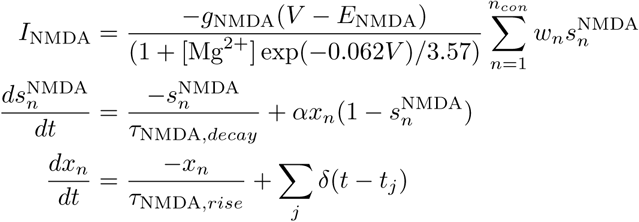

where [Mg^2+^] = 1mM.

Group I mGluRs are postsynaptic and follow the dynamics in Eqs. (1), (2). In neurons with postsynaptic mGluRs, the leak current is decomposed into a scaled K^+^ and non-selective cation leak current (Eq. (3)). The free parameter *τ*_DAG_ differs between CT and TC pathways to match the time course of depolarization observed experimentally [22, 32]. Likewise, *α*_DAG_ is set separately for each pathway to reproduce across-trial changes to thalamic bursting and realistic TC mGluR amplitudes [32, 61]. As described above, trans-thalamic excitatory driver connections possess shortterm synaptic depression and can only activate ionotropic glutamate receptors (AMPA, NMDA). The dynamics of short-term synaptic depression is modeled by the following equation [77]

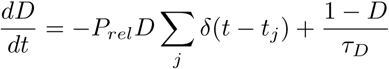

Here, *D* describes the proportion of readily releasable synaptic vesicles and *P_rel_* describes the baseline release probability.

On the other hand, trans-thalamic excitatory modulatory connections possess short-term synaptic facilitation and can activate ionotropic and metabotropic glutamatergic receptors. The dynamics of short-term synaptic facilitation is modeled by the following dynamical equation [77]:

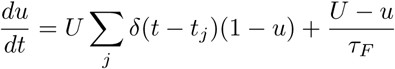

Here, *u* represents the release probability while *U* denotes the baseline for *u*. We note that in facilitating synapses, the term *α*_DAG_ is scaled by *u*.

A complete list of synaptic parameters is provided in Table 3 as well as the associated synaptic weights in Table 4.

**Table 3:** Synaptic Parameters in Trans-thalamic Network.

| Parameter | Value | Reference |
| --- | --- | --- |
| **Synaptic Parameters** |  |  |
| $E_{\text{AMPA}}$ | 0 mV | [17] |
| $E_{\text{NMDA}}$ | 0 mV | [17] |
| $E_{\text{GABA}}$ | -70 mV | [17] |
| $\tau_{\text{AMPA}}$ | 2 ms | [17] |
| $\tau_{\text{NMDA, decay}}$ | 100 ms | [17] |
| $\tau_{\text{NMDA, rise}}$ | 2 ms | [17] |
| $\tau_{\text{GABA, dend}}$ | 20 ms | [54] |
| $\tau_{\text{GABA, soma}}$ | 5 ms | [54] |
| $\alpha_{\text{NMDA}}$ | $0.5 \text{ ms}^{-1}$ | [17] |
| $\beta_K$ | $5.22\text{e-}4 \mu\text{M}^{-1}\text{ms}^{-1}$ | |
| $K_{\text{min}}$ | 0.2 | |
| $E_{\text{NL}}$ | 0 mV | [78] |
| $E_{\text{KL}}$ | -100 mV | |
| $\tau_K$ | 331.6 ms | |
| **Thalamocortical Modulators** |  |  |
| $\tau_{\text{DAG}}$ | 150 ms | [32] |
| **Corticothalamic Modulators** |  |  |
| $\tau_{\text{DAG}}$ | 3000 ms | [22] |
| $\alpha_{\text{DAG}}$ | $0.05 \mu\text{M}$ | |
| **L5 PNs** |  |  |
| $g_{\text{AMPA, soma, ext}}$ | 3.0 nS | |
| $g_{\text{AMPA, dend, ext}}$ | 0.5 nS | |
| $g_{\text{AMPA, soma}}$ | 0.227 nS | |
| $g_{\text{AMPA, dend}}$ | 0.250 nS | |
| $g_{\text{NMDA, soma}}$ | 1.62 nS | |

Table 3 – *Continued from previous page*
| Parameter | Value | Reference |
| --- | --- | --- |
| $g_{\text{NMDA, dend}}$ | 0.7 nS | |
| $g_{\text{GABA, soma}}$ | 12.5 nS | |
| $g_{\text{GABA, dend}}$ | 10.625 nS | |
| **L6 PNs** |  |  |
| $g_{\text{AMPA, ext}}$ | 1.8 nS | |
| $g_{\text{AMPA}}$ | 0.168 nS | |
| $g_{\text{NMDA}}$ | 0.336 nS | |
| $g_{\text{GABA}}$ | 1.5 nS | |
| **Thalamic Neurons** |  |  |
| $g_{\text{AMPA, ext}}$ | 0.32 nS | |
| $g_{\text{AMPA}}$ | 18 nS | [84] |
| $g_{\text{NMDA}}$ | 28 nS | [84] |
| $g_{\text{GABA}}$ | 2.5 nS | |
| **PV INs** |  |  |
| $g_{\text{AMPA, ext}}$ | 0.6 nS | |
| $g_{\text{AMPA}}$ | 0.25 nS | |
| $g_{\text{NMDA}}$ | 0.625 nS | |
| $g_{\text{GABA}}$ | 1.88 nS | |
| **SST INs** |  |  |
| $g_{\text{AMPA, ext}}$ | 0.25 nS | |
| $g_{\text{AMPA}}$ | 0.25 nS | |
| $g_{\text{NMDA}}$ | 0.5 nS | |
| $g_{\text{GABA}}$ | 0.625 nS | |
| **VIP INs** |  |  |
| $g_{\text{AMPA, ext}}$ | 0.18 nS | |
| $g_{\text{AMPA}}$ | 0.1 nS | |
| $g_{\text{NMDA}}$ | 0.12 nS | |
| **TRN INs** |  |  |
| $g_{\text{AMPA, ext}}$ | 1.2 nS | |
| $g_{\text{AMPA}}$ | 0.156 nS | |
| $g_{\text{NMDA}}$ | 0.1875 nS | |
| **Short-Term Synaptic Plasticity** |  |  |
| $\tau_D$ | 423 ms | [45] |
| $\tau_F$ | 250 ms | [30] |
| $P_{\text{rel}}$ | 0.6 | [77] |
| $U$ | 0.1 | [77] |

**Table 4.** Network Connectivity Matrix - Synaptic Weights.

| Pre \ Post | L5 | L6 | Dec | ThalFF | ThalFB | PV | SST | VIP | PV,L6 | PV,D | SST,D | VIP,D | TRN |
| --- | --- | --- | --- | --- | --- | --- | --- | --- | --- | --- | --- | --- | --- |
| L5 | 0.5 | - | 0.4 | 1.0 | 0.9 | 0.75 | 0.1 | - | - | 0.1 | 0.01 | - | 0.3 |
| L6 | - | 0.5 | - | 0.9 | 1.0 | - | - | - | 0.15 | - | - | - | 1.25 |
| Decision ctx | - | - | 0.825 | - | - | 0.2 | 0.01 | - | - | 0.85 | 0.2 | 0.15 | - |
| Thal FF | - | - | 0.3 | - | - | - | - | - | - | - | - | - | - |
| Thal FB | 1.0 | - | - | - | - | - | - | 1.0 | - | - | - | - | - |
| PV | 0.6 | - | - | - | - | 3.0 | - | - | - | - | - | - | - |
| SST | 0.8 | - | - | - | - | 1.0 | - | - | - | - | - | - | - |
| VIP | - | - | - | - | - | - | 1.0 | - | - | - | - | - | - |
| (PV, L6) | - | 0.5 | - | - | - | - | - | - | - | - | - | - | - |
| (PV, decision) | - | - | 0.6 | - | - | - | - | - | - | 2.75 | - | - | - |
| (SST, decision) | - | - | 0.85 | - | - | - | - | - | - | 1.0 | - | - | - |
| (VIP, decision) | - | - | - | - | - | - | - | - | - | - | 1.0 | - | - |
| TRN | - | - | - | 1.0 | 1.0 | - | - | - | - | - | - | - | - |

#### 4.4.3 Task Designs

Prior to all simulations, the network’s spontaneous activity was run for 15 s to allow all slow mGluR-related variables to reach approximate steady state for a given parameterization of *α*_DAG_; the resulting mGluR state at the end of this 15 s period was saved and used to initialize all subsequent trials.

In this work, we employ two task designs to probe the functional consequences of trans-thalamic mGluR modulation. We first mimicked optogenetic activation of layer 6 CT neurons following the protocol of [61]. In our simulation, we delivered trains of brief current pulses (235 pA, 2 ms) to L6 neurons at 10 Hz for 1 s. Each train was preceded by a 500 ms pre-stimulus period and followed by a 2 s ITI.

We next had the network perform a detection task where a stimulus was delivered to the sensory area (L5 and L6 neurons) through external AMPA synapses following a fixation period of 500 ms. The stimulus consisted of Poisson spike trains with time-dependent rate *r*(*t*), applied for 150 ms. During stimulus presentation, *r*(*t*) was drawn from a Gaussian distribution with mean *µ* and standard deviation *σ* (8 Hz). Network decisions were read out following stimulus offset by determining whether the decision area exhibited sustained activity, defined as a firing rate exceeding 20 Hz for at least 200 ms. A trial in which the decision area lacked sustained activity was classified as a miss whereas if the decision area generated sustained activity the trial was classified as a hit. To generate psychometric functions, we varied the input strength *µ* between 49 and 66 Hz (which were subsequently normalized from 0 to 1, for visualization purposes). For each value of *µ*, the average performance of the network in the detection task was calculated from 135 independent stimulus presentations. Finally, to quantify history-dependent effects, we introduced a variable-length ITI (default: 2 s) where stimuli of a fixed input strength were presented in sequences of three consecutive trials; an independent sequence was repeated 135 times and the average performance in each trial across the sequences was computed.

## Acknowledgements.

thank Vincent Huson and Wade Regehr for generously providing mGluR EPSC traces from cerebellar unipolar brush cells, and Megan Kirchgessner and Ed Callaway for sharing *in vivo* extracellular pulvinar recordings and analysis code. We thank John Rinzel, Jorge Jaramillo, Federico Brandalise, Anthony Holtmaat, and all members of the Wang Lab for their helpful comments on this work. This project was partly supported by NEURONEX (NSF 2015276) and a T32 NIMH Training Fellowship in Learning, Memory, Development, and Plasticity (to AL).

---

**Fig. S1.**
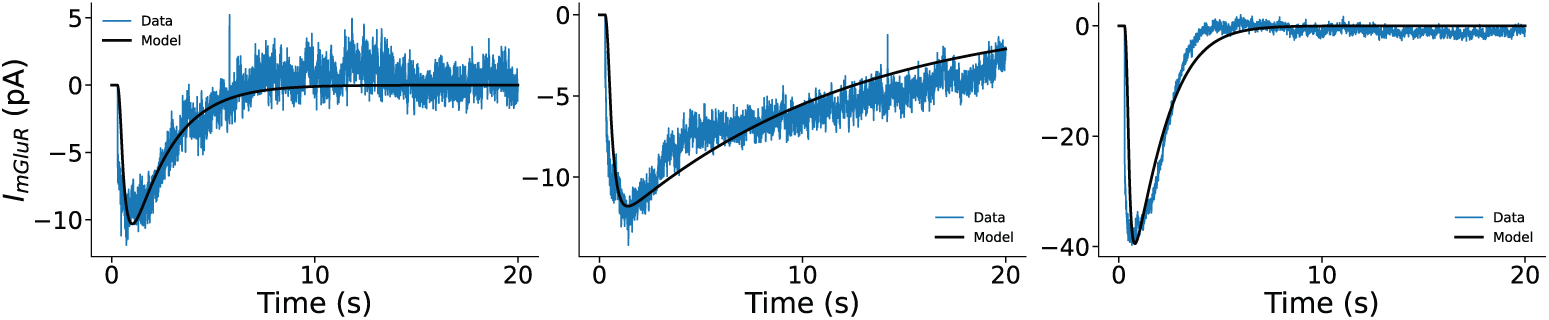
Additional example fits. Three example mGluR EPSCs fit with *α*_DAG_ = 0.0997 *µ*M, *τ*_DAG_ = 1820 ms (left), *α*_DAG_ = 0.09067 *µ*M, *τ*_DAG_ = 10000 ms (middle), and *α*_DAG_ = 0.5711 *µ*M, *τ*_DAG_ = 1166.03 ms (right).

**Fig. S2.**
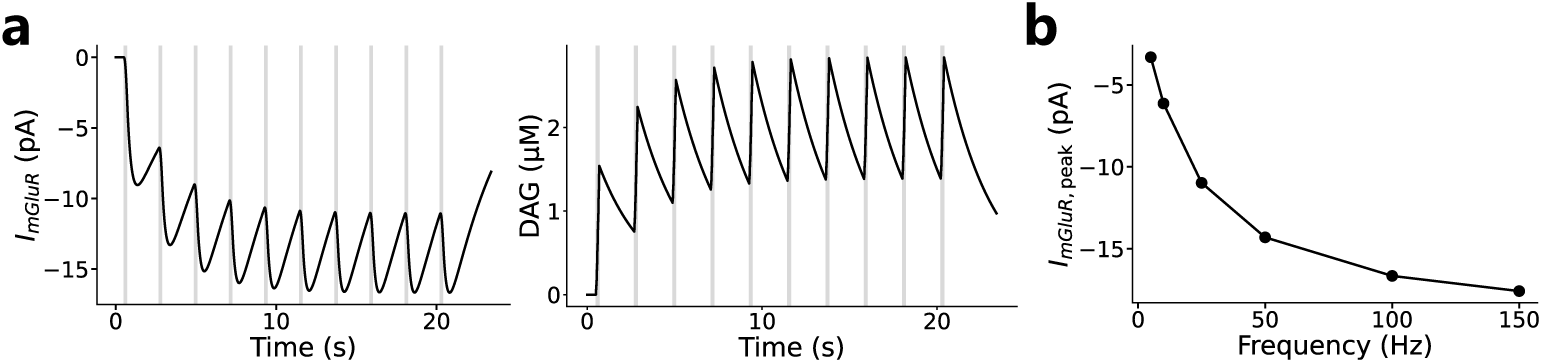
Slow Kinetics of mGluR in the presence of short-term synaptic facilitation. An example synapse onto model cerebellar unipolar brush cells is endowed with short-term synaptic facilitation dynamics. **(a.)** Even in the presence of facilitation, with *α*_DAG_ = 0.135 *µ*M*, τ*_DAG_ = 2800 ms, the model displays accumulation of mGluR EPSCs resulting from presynaptic input (20 × 100 Hz, time period denoted by gray bars) due to DAG’s slow decay. **(b.)** The saturated peak mGluR EPSC as a function of input frequency has an equivalent relationship as what is seen in Fig. 2.

**Fig. S3.**
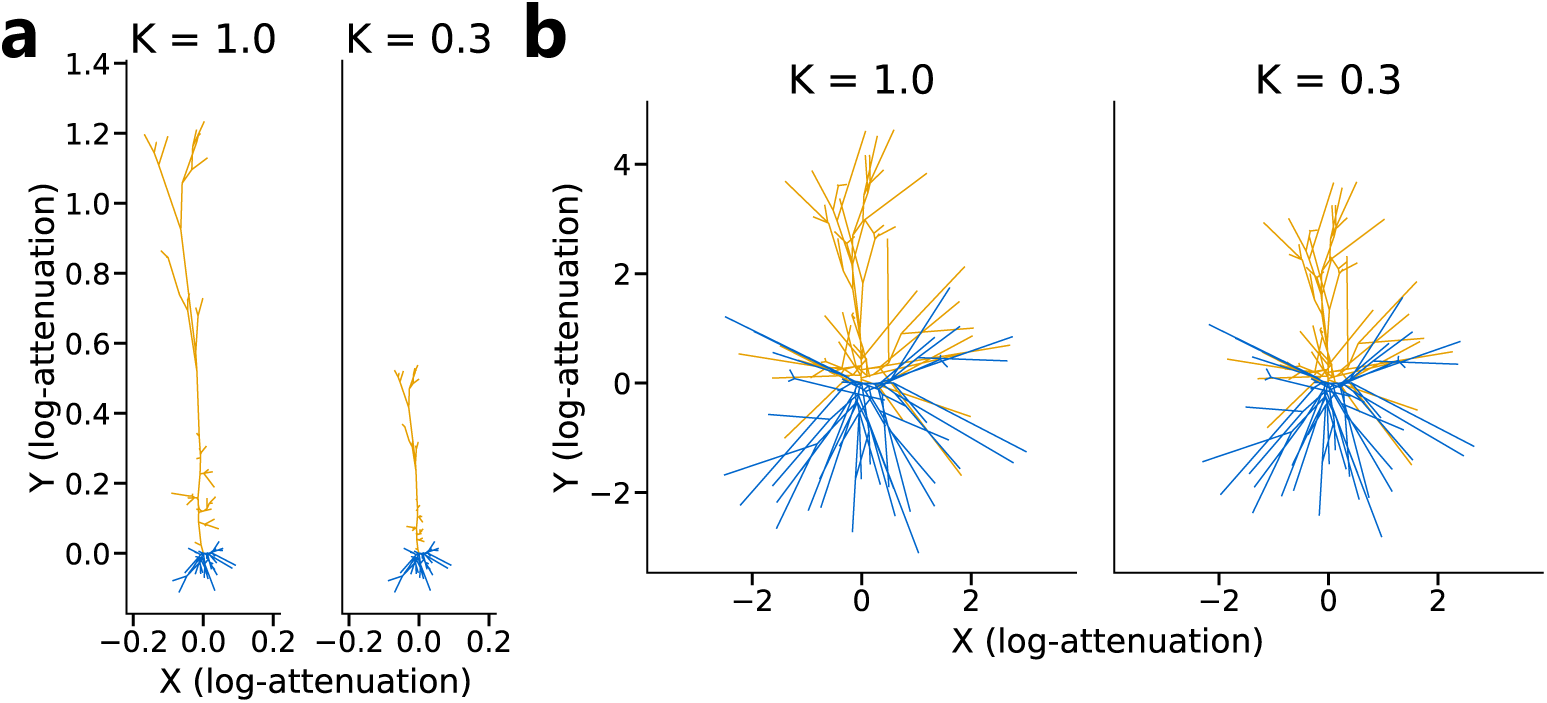
Functional attenograms of L5 PN following K2P suppression. **(a.)** Centrifugal somatic attenogram. Distance in this space corresponds to the log attenuation of voltage propagating from the soma toward distal dendrites (see Methods for details). From left to right, increasing mGluR-mediated suppression of K2Ps in apical dendritic compartments reduces attenuation, indicating that somatic depolarizations propagate more effectively into distal dendrites. **(b.)** Centripetal somatic attenogram. Distance in this space corresponds to the log attenuation of voltage propagating from the dendrites toward the soma. From left to right, increasing mGluR-mediated suppression of K2Ps in apical dendritic compartments reduces attenuation, reflecting stronger dendritic integration and electrotonic shortening of distal apical branches to the soma.

**Fig. S4.**
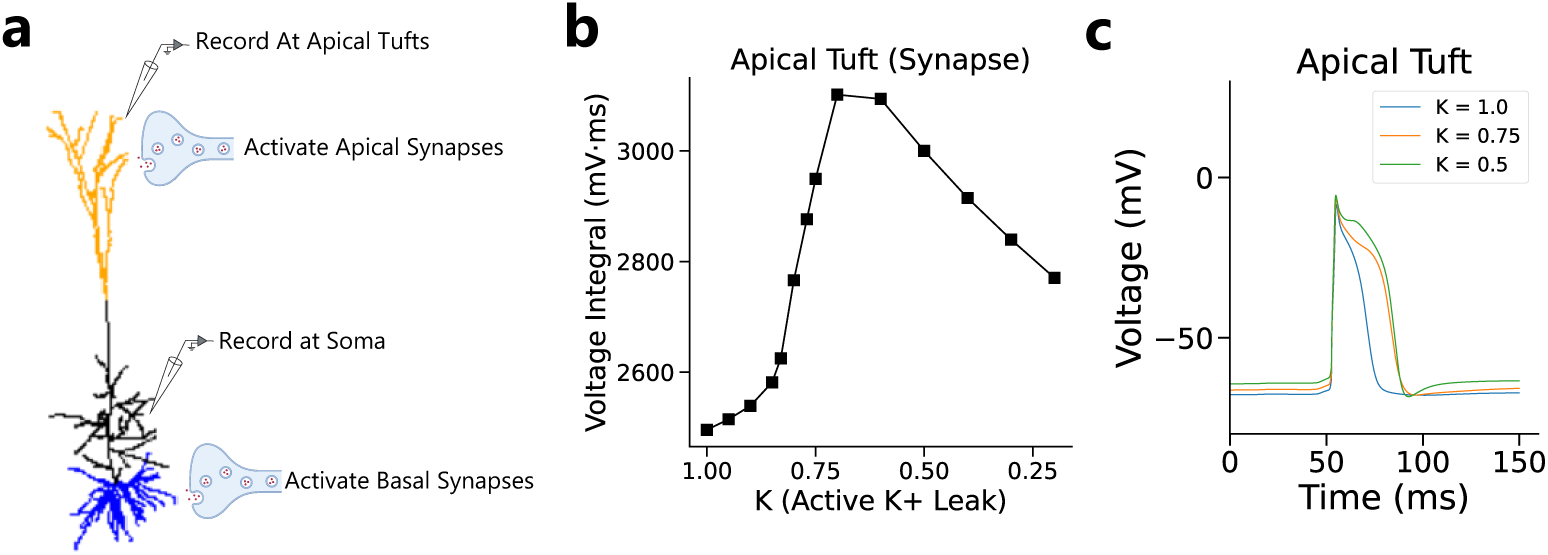
K2P suppression enhances dendritic NMDA plateau potentials. **(a.)** Experimental protocol in multicompartment model. A fixed coincident synaptic input is provided to synapses at apical and basal dendrites (11 tuft and 9 basal inputs) while simultaneously recording the membrane voltage at activated apical tufts and the soma. **(b.)** Voltage integrals changed non-linearly as *K* decreased, revealing a sharp increase in the dendritic voltage integral with sufficiently small *K* (*K <* 0.8) coinciding with the onset of a complete dendritic NMDA spike waveform. **(c.)** Example voltage traces at the apical tuft display an NMDA spike with sufficient mGluR-mediated suppression of K2P channels.

**Fig. S5.**
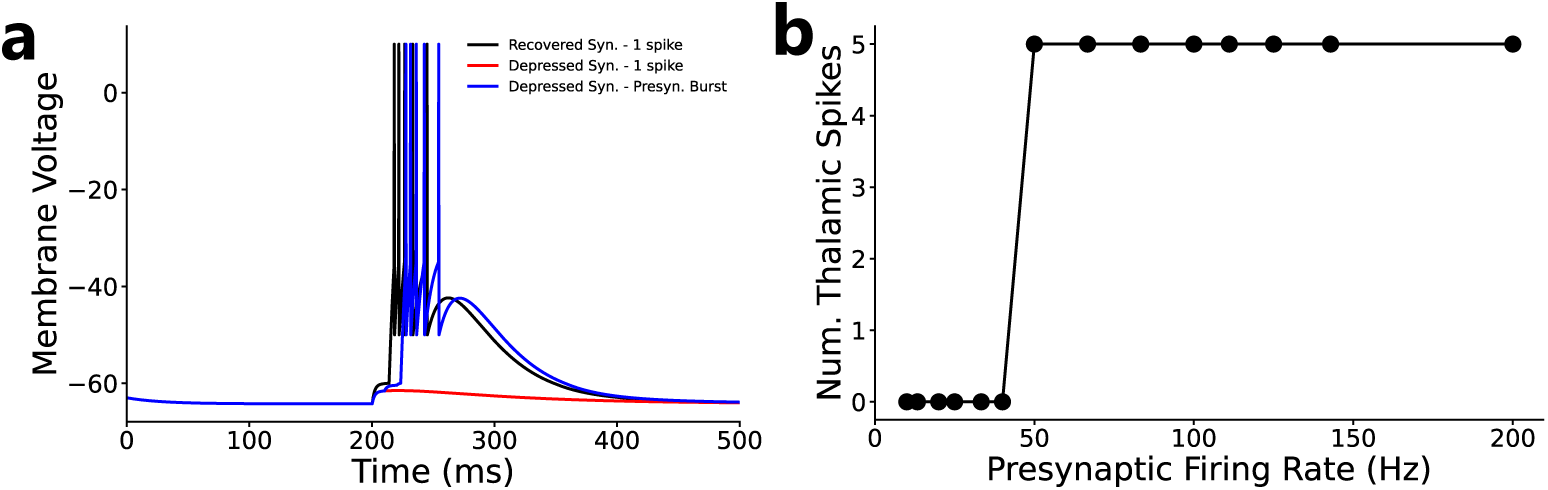
L5 Driver Synapses Requires High Presynaptic Firing Rates to Drive Thalamic Responses when synapses are Depressed. **(a.)** Simulated thalamic membrane voltage in response to L5 cortical input under three conditions: a fully recovered synapse receiving a single spike (black), a depressed synapse receiving a single spike (red), and a depressed synapse receiving a burst of 3 spikes at 100 Hz (blue). While a single input through a recovered synapse is sufficient to evoke a thalamic action potential, synaptic depression causes depolarization resulting from single spikes to be subthreshold. Only when presynaptic spikes arrive in rapid succession does the depressed synapse reliably drive thalamic firing. **(b.)** Number of thalamic spikes evoked as a function of presynaptic firing rate, for a fixed number of pulses (3 presynaptic events). Thalamic spiking is absent at low firing rates and emerges sharply only above a critical rate threshold, illustrating that high-frequency cortical activity is necessary to overcome synaptic depression and reliably recruit thalamic responses in L5 driver synapses.

**Fig. S6.**
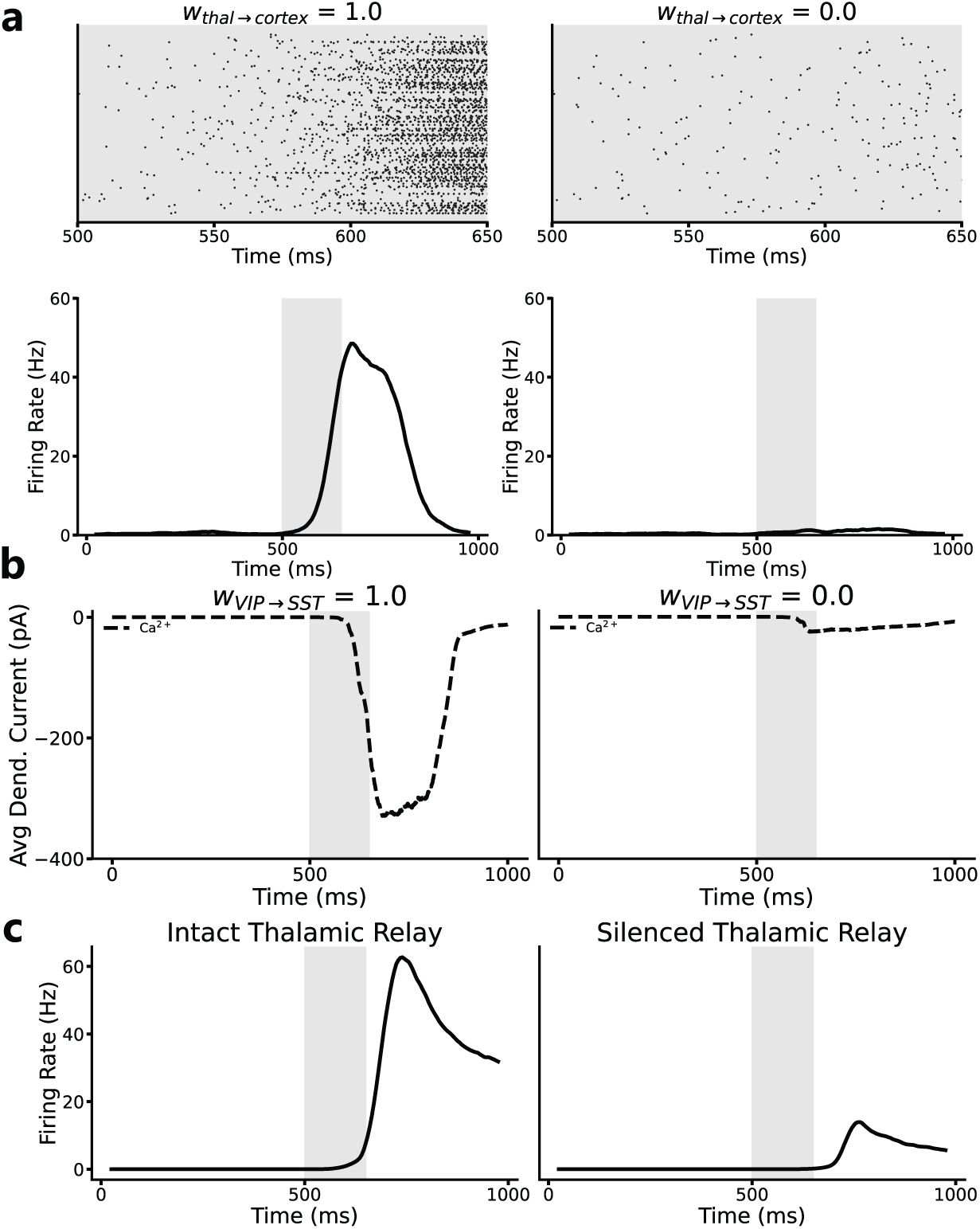
Network Design Constraints. (**a.**) Raster plots of sensory-area neurons (top) and their population firing rates (bottom) with TC feedback onto sensory cortex intact (*w*_thal_*_→_*_cortex_ = 1.0) versus removed (*w*_thal_*_→_*_cortex_ = 0.0). Without thalamic feedback, stimulus-driven cortical activity is substantially reduced, consistent with [52]. (**b.**) Average dendritic Ca^2+^ currents in sensory-area PNs with the VIP→SST disinhibitory motif intact (*w*_VIP_*_→_*_SST_ = 1.0) versus removed (*w*_VIP_*_→_*_SST_ = 0.0). Recruited by thalamic feedback via VIP interneurons, this motif gates dendritic Ca^2+^ plateau potentials by opening temporally precise windows of reduced SST-mediated dendritic inhibition [53]. (**c.**) Decision-area population firing rate with the higher-order sensory thalamic relay to the decision area intact (left) versus silenced (right); silencing the relay markedly reduces the stimulus-evoked decision-area response with the network relying solely on cortico-cortical connectivity. Gray bars mark stimulus onset. Rasters show a single representative trial; firing-rate and dendritic-current traces are averaged over 10 trials of medium-strength stimulus presentation.

**Fig. S7.**
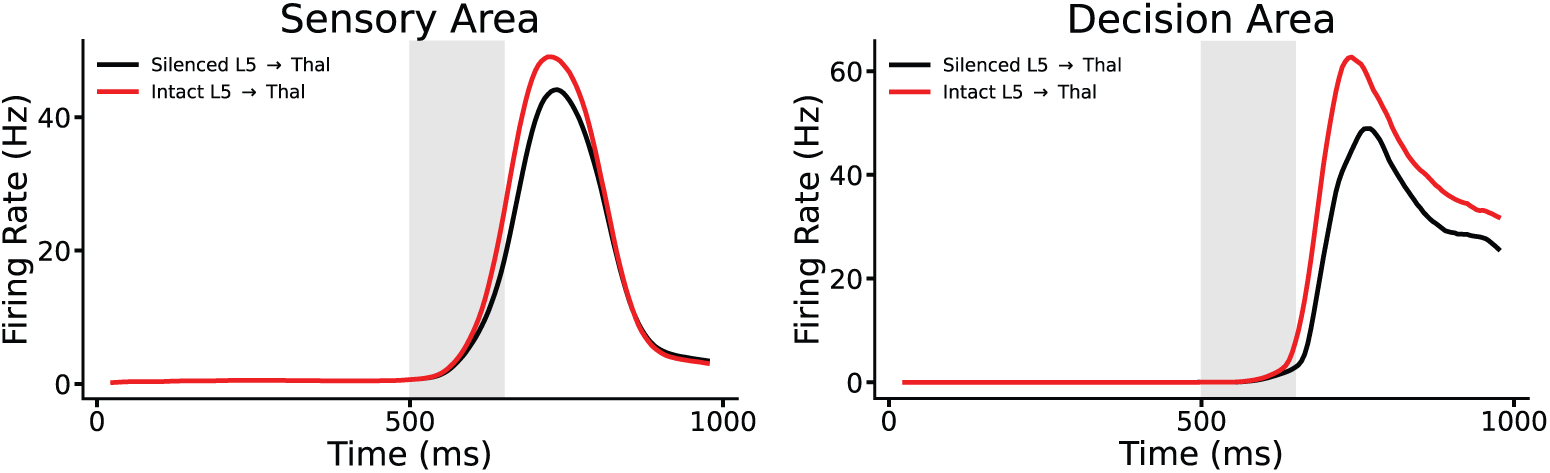
Silencing L5 to HOs thalamus projection attenuates perceptual detection. Silencing L5 to HOs thalamus projections during stimulus presentation (black) reduces stimulus-evoked firing in both cortical areas. In the sensory area (left), this reflects the loss of depolarizing L5 CT input, which weakens TC modulation of sensory cortex. In the decision area (right), firing is further reduced because silencing this projection also removes the thalamic relay that links the two cortical regions, lowering effective connectivity between them.

**Fig. S8.**
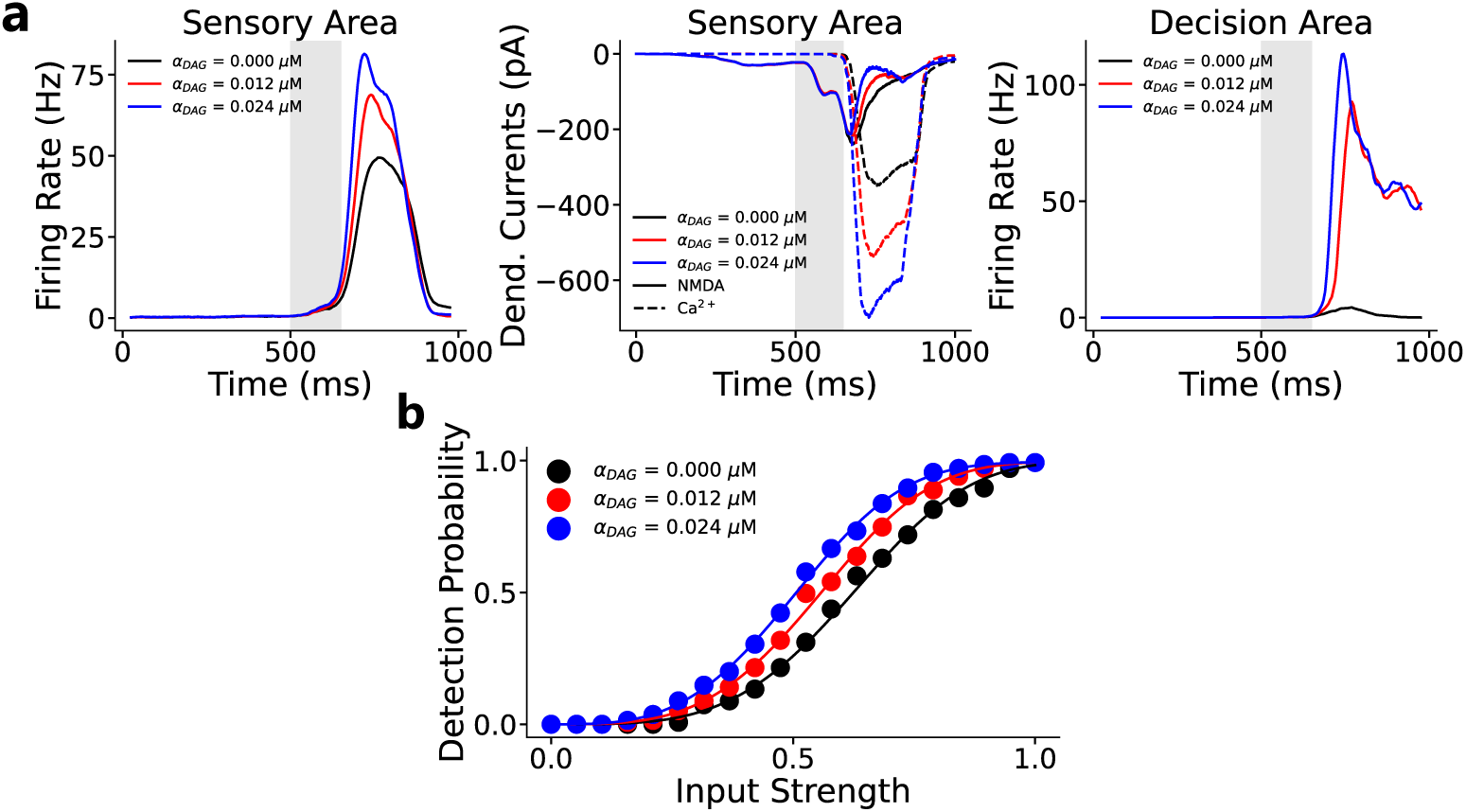
TC mGluR Activation Lowers Perceptual Threshold in the Tonic Firing Regime. We repeated the simulations of Fig. 4 with a 30 pA constant current injection to HOs thalamic neurons (in addition to their baseline CT mGluR depolarization), causing them to fire tonically rather than in burst mode. **(a.)** An example trial showing sensory area firing rates (left), sensory area dendritic currents (middle), and decision area firing rates (right) when we increase mGluR activation (*α*_DAG_ = 0.000, 0.012, 0.024 *µ*M). As was the case in Fig. 4, stronger mGluR engagement increases the firing of sensory neurons, reflected in larger voltage-dependent Ca^2+^ (L-type) currents, and drives the decision area toward sustained activity. **(b.)** Detection probability as a function of input strength for the three levels of mGluR activation. Increasing *α*_DAG_ shifts the psychometric curve leftward, lowering the perceptual threshold.

**Fig. S9.**
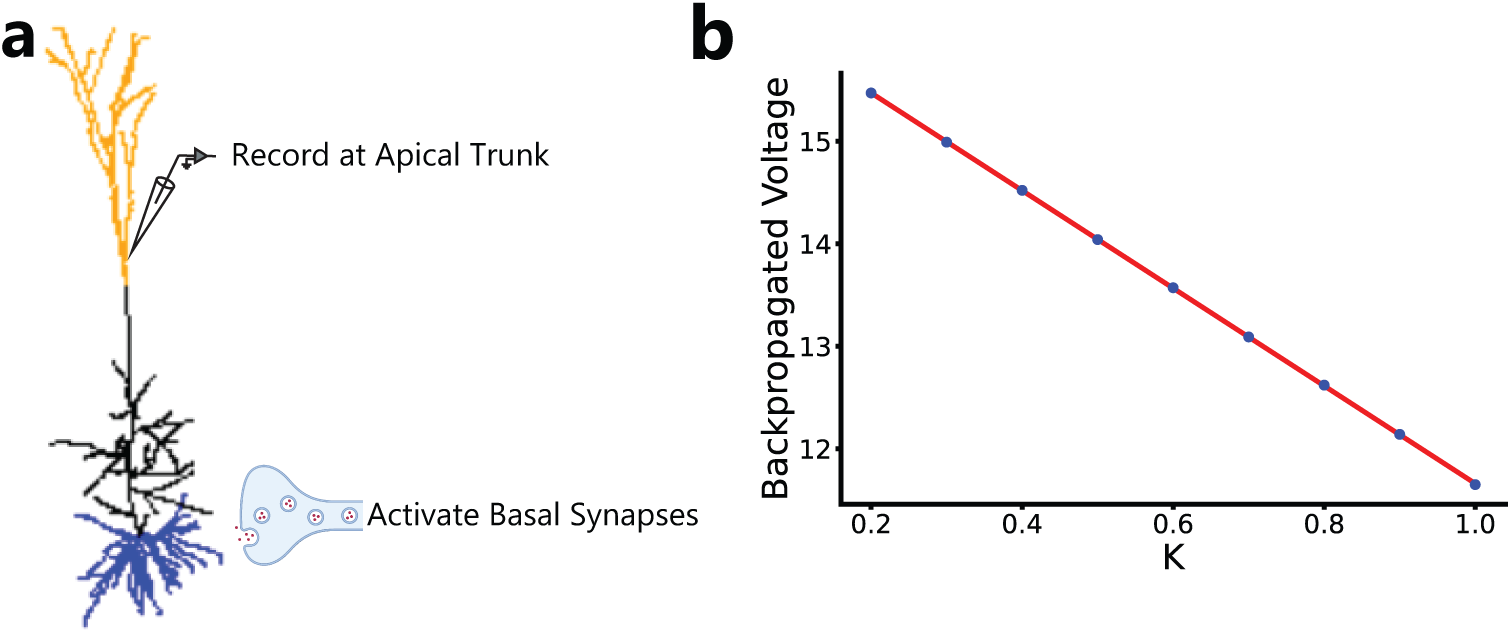
Backpropagating action potential calibration. (**a.**) Schematic of the multicompartmental PN model used to calibrate the bAP in two-compartment PNs in the spiking network. Basal synapses are activated to elicit a somatic spike, and the resulting backpropagated voltage is recorded at the apical trunk. (**b.**) Relationship between the fraction of open K2P channels (*K*) and bAP amplitude measured at the apical trunk. A linear fit ((16.431 − 4.783*K*) mV, red) captures the *K*-dependent voltage response used to set the bAP magnitude in the reduced two-compartment model.

